# The Impact of TOR-S6K-eIF5A Signaling on Translational Control is Key to Plant Root Development

**DOI:** 10.64898/2026.09.16.752225

**Authors:** Joong-Tak Yoon, Du-Hwa Lee, Sejin Choi, Chang Sook Ahn, Shaban Muhammad, Yicheng Liu, Hyun-Sook Pai, Eun Yu Kim, Ho-Seok Lee

## Abstract

Nutrient-responsive Target of Rapamycin (TOR)-S6 kinase (S6K) signaling coordinates plant growth with protein synthesis, but how it interfaces with translation elongation to produce specific developmental outputs remains unclear. Here, we identify eukaryotic translation factor 5A (eIF5A) as an S6K-associated phosphoprotein in *Arabidopsis thaliana*. S6K1 and S6K2 associate with all three eIF5A isoforms and phosphorylate them in vitro, with Ser2 emerging as the major S6K1-responsive site in eIF5A-2. The corresponding N-terminal serine is invariant across the analyzed Archaeplastida eIF5A proteins. Conditional depletion of TOR, S6K1/2, or eIF5A produces overlapping reductions in primary root length and root hair coverage. eIF5A depletion leaves bulk protein synthesis and polysome profiles largely unchanged while selectively decreasing output from deca-proline reporters. Reporter activity is restored by amiRNA-resistant wild-type and phosphomimetic S2D eIF5A-2, whereas S2A does not restore activity, demonstrating the functional importance of the Ser2 state. Proteomic profiling identifies a restricted set of eIF5A-responsive proteins, and five of six tested insertion mutants display altered primary root growth, root hair coverage, or both. Together, these findings uncover a regulatory connection between S6K and eIF5A and establish eIF5A-dependent selective translation as a mechanism contributing to root development in *A. thaliana*.

## Introduction

Cell growth and development fundamentally depend on the precise regulation of protein synthesis(Liu & Sabatini, 2020; Saxton & Sabatini, 2017; Scarpin et al., 2022). In eukaryotes, the Target of Rapamycin (TOR) signaling pathway functions as a central hub that integrates nutrient availability and environmental cues to coordinate anabolic processes, particularly translation(Burkart & Brandizzi, 2021; Jewell et al., 2013; Liu & Sabatini, 2020; Saxton & Sabatini, 2017; Scarpin et al., 2022). This pathway is evolutionarily conserved across eukaryotes, including yeast, mammals, and plants, ensuring that cellular growth is tightly coupled to metabolic status(Ingargiola et al., 2020; Loewith & Hall, 2011; Menand et al., 2002; Saxton & Sabatini, 2017; Scarpin et al., 2022; Shi et al., 2018; Wullschleger et al., 2006; Xiong & Sheen, 2014). In plants, glucose-TOR signaling activates root meristems and promotes root growth in response to nutrient availability(Dong et al., 2022; Li et al., 2017; Xiong et al., 2013; Xiong & Sheen, 2012). However, the molecular mechanisms by which TOR-dependent translational regulation shapes these developmental outputs remain unclear(Chen et al., 2018; Scarpin et al., 2020; Scarpin et al., 2022).

Major downstream effectors of TOR include S6 Kinase 1 and 2 (S6K1/2), which modulate translational capacity by phosphorylating components of the translational machinery, including ribosomal protein S6 (RPS6)(Chen et al., 2018; Mahfouz et al., 2006; Turck et al., 2004). However, plant S6Ks are not merely general amplifiers of TOR-dependent protein synthesis. S6K1 also phosphorylates eukaryotic translation initiation factor 3h (eIF3h) to stimulate translation reinitiation of selected uORF-containing mRNAs(Schepetilnikov et al., 2013). This provides a direct example of transcript-selective translational control downstream of TOR-S6K signaling. Consistently, translatome-wide analyses have shown that TOR perturbation does not affect all mRNAs uniformly, but preferentially remodels the translation of defined transcript classes, including 5’TOP-containing mRNAs bound by the TOR effector La-related protein 1 (LARP1)(Scarpin et al., 2020). These findings indicate that TOR-dependent translational output is specified through downstream effectors that act at both global and transcript-selective levels(Scarpin et al., 2020; Scarpin et al., 2022; Schepetilnikov et al., 2013). While the role of S6K in conferring translational specificity is increasingly recognized, how this selectivity is converted into specific developmental outputs remains unclear.

Eukaryotic initiation factor 5A (eIF5A) is a conserved translation factor that promotes elongation at multiple difficult-to-translate sequence contexts and also contributes to termination(Gutierrez et al., 2013; Pelechano & Alepuz, 2017; Schuller & Green, 2018; Schuller et al., 2017). During translation elongation, the ribosome sequentially catalyzes peptide bond formation at the peptidyl transferase center. However, this process is not uniform. Proline is incorporated more slowly than alanine or phenylalanine because its N-alkylated cyclic structure creates unfavorable steric and chemical constraints during accommodation and peptidyl transfer(Pavlov et al., 2009). Stalling is especially prominent in some polyproline and XPPX contexts, but its severity depends strongly on the surrounding sequence(Peil et al., 2013; Starosta et al., 2014). Structural work attributes polyproline stalling to unfavorable positioning and dynamics of the peptidyl-transfer substrates and shows how EF-P can restore a productive geometry(Huter et al., 2017). This activity has been well characterized for eIF5A in yeast and mammals and for its bacterial functional analog EF-P(Barba-Aliaga et al., 2021; Doerfel et al., 2013; Gutierrez et al., 2013; Huter et al., 2017; Saini et al., 2009; Schuller et al., 2017; Ude et al., 2013). A defining feature of eIF5A is hypusination, a unique post-translational modification of a conserved lysine generated through the sequential actions of deoxyhypusine synthase (DHS) and deoxyhypusine hydroxylase (DOHH)(Park et al., 2006; Park & Wolff, 2018). Hypusination strongly enhances eIF5A activity in ribosome-dependent translation assays(Park et al., 2011; Saini et al., 2009). In *Arabidopsis thaliana*, eIF5A hypusination responds to abscisic acid, and DHS suppression implicates the pathway in root architecture, root hair development, senescence, bolting, and other developmental traits(Belda-Palazon et al., 2016; Belda-Palazon et al., 2014; Palfi et al., 2021; Wang et al., 2003). In plants, eIF5A isoforms have been implicated in growth, development, and stress responses, including cytokinin-mediated protoxylem differentiation and programmed cell death during pathogen infection(Feng et al., 2007; Hopkins et al., 2008; Ma et al., 2010; Ren et al., 2013). Beyond hypusination, plant eIF5A is phosphorylated at its N-terminal Ser2. CK2 phosphorylates maize eIF5A at Ser2(Łebska et al., 2010; Lewandowska-Gnatowska et al., 2011), and *A. thaliana* phosphoproteomic data associate eIF5A-2/eIF5A-3 Ser2 phosphorylation with light and cellular energy status(Boex-Fontvieille et al., 2013; Nukarinen et al., 2016). However, whether plant eIF5A promotes translation through such elongation-sensitive sequences and how its activity is connected to nutrient-responsive signaling remains unresolved.

Here, we identify *A. thaliana* eIF5A as an S6K-associated phosphoprotein and establish its N-terminal Ser2 as a conserved S6K1-responsive site. S6K1 and S6K2 associate with all three eIF5A isoforms and phosphorylate them in vitro, whereas conditional depletion of TOR, S6K1/2, or eIF5A produces overlapping reductions in primary root growth and root hair coverage. eIF5A depletion selectively reduces translation through polyproline sequences without measurably altering bulk protein synthesis or polysome profiles, and reporter rescue by wild-type and S2D, but not S2A, eIF5A-2 demonstrates the functional importance of the Ser2 state. Proteomic and genetic analyses further identify eIF5A-responsive proteins that support root development. Together, these findings uncover an S6K-eIF5A regulatory connection and establish eIF5A-dependent selective translation as a mechanism contributing to root development in *A. thaliana*.

## Results

### TOR-S6K signaling regulates root growth and root hair coverage

TOR kinase is required for root growth and root hair formation (Montané & Menand, 2013; Xiong & Sheen, 2012), but whether S6Ks mediated these effects has not been established genetically. To address this, we analyzed the phenotypes of estradiol-inducible *TOR* RNAi lines (*tor-es #1* and *tor-es #2*) (Xiong & Sheen, 2012) and estradiol-inducible artificial microRNA (amiRNA) lines targeting both *S6K1* and *S6K2* (*s6k1/2-es #2* and *s6k1/2-es #11*). Both *TOR* and *S6K1/2*-silenced lines treated with 10 μM β-estradiol (Es) showed impaired root growth and root hair elongation compared to the EtOH control (Figures 1A and S1). To quantify these phenotypes, we measured primary root length and root hair coverage. Root hair coverage was calculated as the ratio of the root hair-occupied region to the total primary root length, enabling normalization of root hair development to overall root growth and distinguishing root hair-specific defects from general growth reduction caused by TOR signaling perturbation (Figure 1B). Quantitative analysis revealed significant reductions in both root length and root hair coverage in Es-treated *TOR* and *S6K1/2*-silenced lines (Figures 1C and 1D). These results indicate that TOR promotes primary root growth and root hair elongation through S6K signaling.

**Figure 1.**
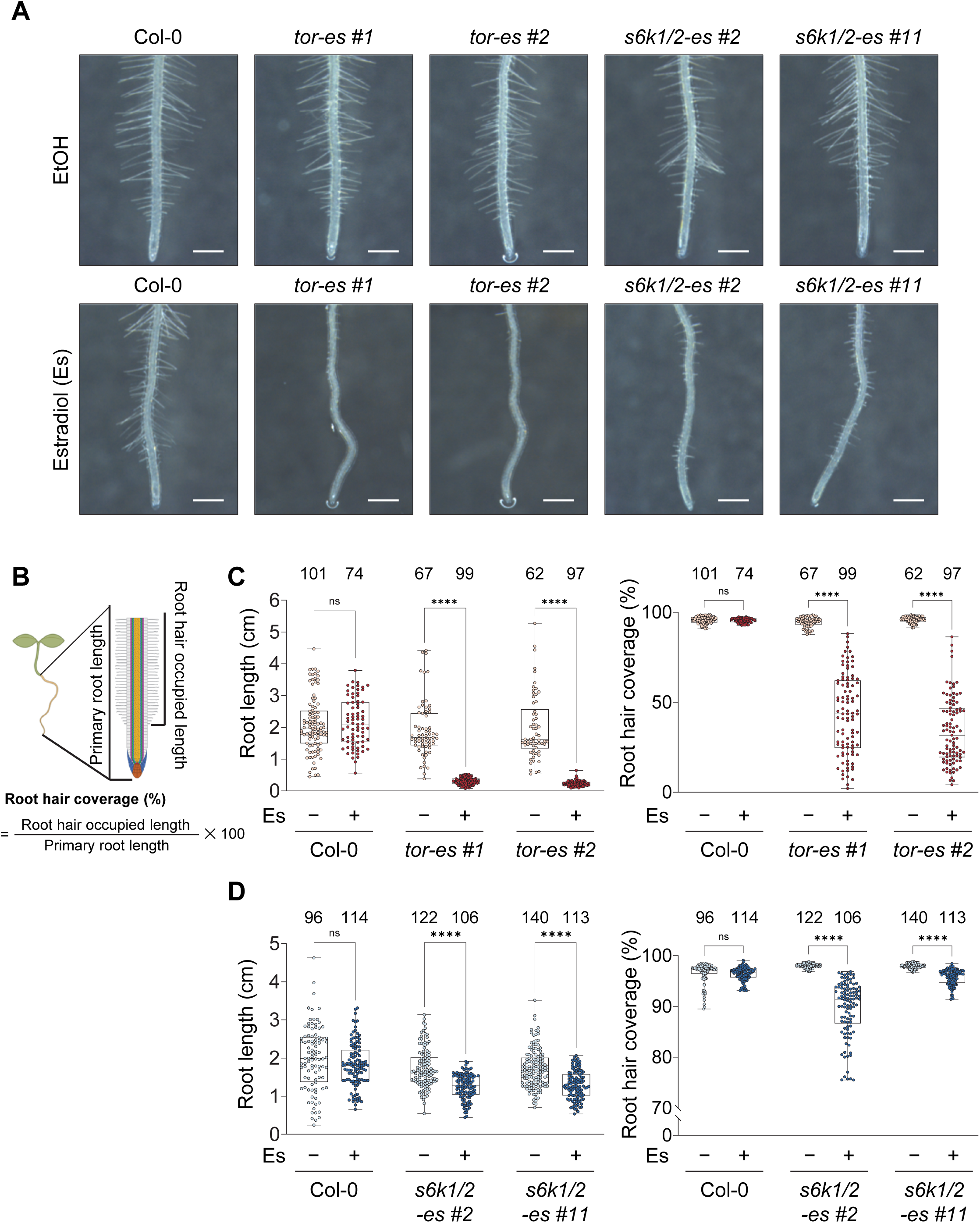
TOR and S6K are required for primary root growth and root hair formation. (A) Representative root images of Col-0, estradiol-inducible *TOR* RNAi (*tor-es #1* and *tor-es #2*), and estradiol-inducible *S6K1/2* amiRNA (*s6k1/2-es #2* and *s6k1/2-es #11*) under EtOH or 10 μM estradiol (Es). Scale bars, 1 mm. (B) Schematic illustration showing measurement of primary root length and root hair coverage. Root hair coverage (%) was calculated as root hair-occupied length divided by primary root length. (C and D) Box plots represent quantification of primary root length and root hair coverage in (C) Col-0, *tor-es #1*, and *tor-es #2*, and in (D) Col-0, *s6k1/2-es #2*, and *s6k1/2-es #11* seedlings under EtOH or 10 μM Es. Boxes indicate the first and third quartiles, the center lines indicate the median, and whiskers indicate the minimum and maximum data points. Sample sizes (*n*) for each group are indicated above each plot. Individual data points are shown as dots. Statistical significance was assessed using multiple Welch-corrected unpaired t-tests with Holm–Šídák correction for multiple comparisons to compare EtOH and Es within each genotype; ns, P > 0.05; ****, P < 0.0001.

### eIF5A Ser2 is a conserved S6K phosphorylation site

To identify downstream effectors of S6Ks, we performed immunoprecipitation (IP) coupled with liquid chromatography-tandem mass spectrometry (LC-MS/MS) using 7-day-old *Arabidopsis thaliana* seedlings initially grown under light and glucose conditions (Figure 2A). Col-0 seedlings grown under the same 7-day light and glucose conditions, without subsequent dark and starvation treatment, were harvested and served as a negative control (CTL) for non-specific binding in the immunoprecipitation. The *35S::Flag-S6K1* transgenic line, after the same 7-day light and glucose treatment, was subjected to 24 hours of darkness and starvation and harvested as S6K1-DS. After re-exposure to light and glucose for 6 hours, seedlings were harvested as S6K1-LG. Harvested samples were subjected to Flag affinity purification and analyzed by LC-MS/MS. This analysis identified four candidate phosphopeptides as S6K-associated substrates (Figures 2B). To prioritize candidates potentially relevant to root development, we examined expression data from the *A. thaliana* eFP browser and found that only *AT1G26630*, which encodes eukaryotic initiation factor 5A-2 (eIF5A-2), was highly expressed in roots (Figures S2A-S2D)(Winter et al., 2007). The LC-MS/MS data further showed that eIF5A-2 is phosphorylated at Ser2 (Figure 2C), suggesting that eIF5A-2 is a prominent candidate downstream target of S6K1 relevant to root development.

**Figure 2.**
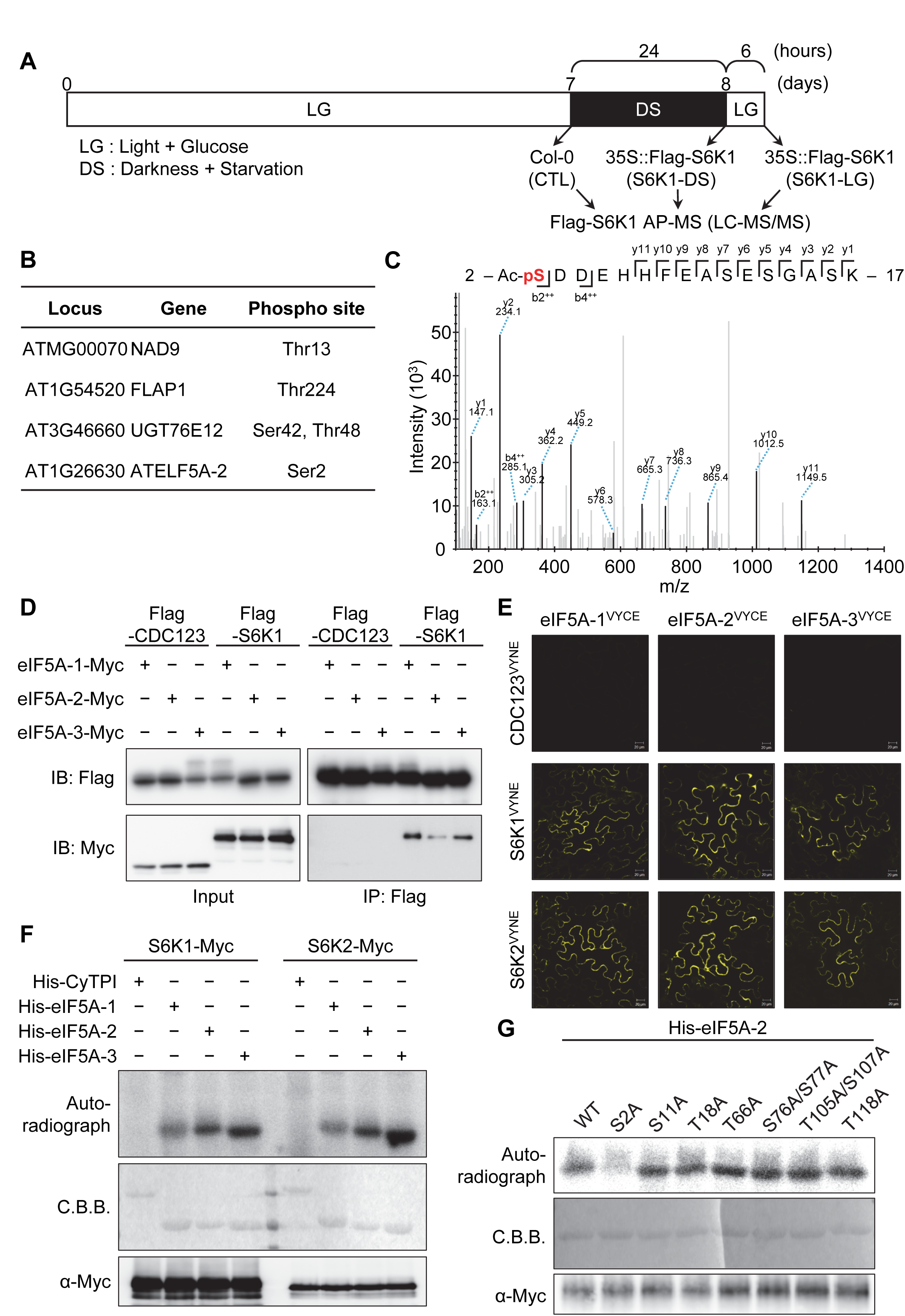
S6K associates with eIF5A and phosphorylates eIF5A-2 at Ser2. (A) Experimental workflow for Flag-S6K1 affinity purification coupled with LC-MS/MS analysis. Col-0 (CTL) and *35S::Flag-S6K1* seedlings were grown for 7 days after germination (DAG) under light and glucose (LG) conditions. CTL samples were collected at 7 DAG and used as a negative control for background subtraction. *35S::Flag-S6K1* seedlings were then transferred to darkness and starvation (DS) conditions for 24h. Samples were collected immediately after DS treatment (S6K1-DS) or after an additional 6 h recovery under LG conditions (S6K1-LG). Flag affinity purification was performed on all samples prior to LC-MS/MS analysis. (B) S6K1-associated phosphoprotein candidates detected from S6K1-LG. Table includes locus, gene names, and phosphorylation site for the four candidates analyzed. (C) LC-MS/MS spectrum of the N-terminal eIF5A-2 phosphopeptide corresponding to residues 2-17 (Ac-pSDDEHHFEASESGASK). The matched *b-* and *y-*ion series support assignment of the phosphorylation site to Ser2. (D) Co-immunoprecipitation (IP) analysis of Flag-CDC123 or Flag-S6K1 with eIF5A-1-Myc, eIF5A-2-Myc, or eIF5A-3-Myc. Flag-CDC123 or Flag-S6K1 was transiently coexpressed with each eIF5A-Myc isoform in *Nicotiana benthamiana* leaves by agroinfiltration. Total protein extracts were subjected to IP with anti-Flag beads and analyzed by immunoblotting using anti-Flag and anti-Myc antibodies. Flag-CDC123 served as a negative control. (E) Bimolecular fluorescence complementation (BiFC) assay showing the interactions of S6K1 or S6K2 with eIF5A isoforms in *N. benthamiana* leaves. S6K1-VYNE or S6K2-VYNE was transiently coexpressed with eIF5A-1-VYCE, eIF5A-2-VYCE, or eIF5A-3-VYCE by agroinfiltration. CDC123-VYNE served as a negative control. YFP fluorescence was visualized by confocal microscopy. Scale bars, 20 μm. VYNE and VYCE represent the N- and C-terminal halves of Venus, respectively. (F and G) In vitro kinase assays showing phosphorylation of His-eIF5A isoforms (F) or His-eIF5A-2 alanine-substituted variants (G) by immunoprecipitated S6K1/2-Myc in the presence of [γ-^32^P]ATP. (F) Immunoprecipitated S6K1-Myc or S6K2-Myc was incubated with recombinant His-CyTPI, His-eIF5A-1, His-eIF5A-2, or His-eIF5A-3. His-CyTPI served as a negative control substrate. (G) Immunoprecipitated S6K1-Myc was incubated with recombinant His-eIF5A-2 WT or the indicated alanine-substituted variants (S2A, S11A, T18A, T66A, S76A/S77A, T105A/S107A, and T118A) to identify the S6K1 phosphorylation site. Phosphorylated substrates were detected by autoradiography. Coomassie Brilliant Blue (CBB) staining shows the recombinant substrate input, and anti-Myc immunoblotting shows the kinase input. CyTPI, cytosolic triose phosphate isomerase.

Given that *A. thaliana* possesses three eIF5A isoforms, we investigated whether S6K proteins could associate with all three eIF5A isoforms in vivo. Accordingly, we conducted co-IP and bimolecular fluorescence complementation (BiFC) assays with a negative control, CDC123 (AT4G05440), an ATP-dependent chaperone involved in eIF2 complex assembly that was not identified in our IP-LC-MS/MS analysis(Chen et al., 2023; Panvert et al., 2015; Perzlmaier et al., 2013). Both assays confirmed that S6K1 and S6K2 interact with all three eIF5A isoforms, but not with CDC123, in the cytosol in vivo (Figures 2D and 2E). We generated an anti-eIF5A antibody against a recombinant His-eIF5A-2 protein and confirmed that it recognizes all three eIF5A isoforms (Figure S3A). Using this antibody, we then performed semi-native co-IP in the *35S::Flag-S6K1 A. thaliana* line. Endogenous eIF5A proteins were immunoprecipitated with the anti-eIF5A antibody, and co-precipitated Flag-S6K1 was detected by immunoblotting, confirming the association between S6K1 and endogenous eIF5A isoforms. Notably, glucose treatment increased the amount of co-precipitated Flag-S6K1, with the appearance of a phosphorylated S6K1 band, suggesting that the S6K1-eIF5A association is enhanced upon glucose-induced S6K activation (Figure S3B). Together, these results extend the IP-LC-MS/MS-based identification of eIF5A-2 to the eIF5A family and demonstrate a physical interaction between S6K and eIF5A in vivo.

To determine whether eIF5A is a direct substrate of S6K, we performed an in vitro kinase assay using recombinant His-tagged eIF5A isoforms and immunoprecipitated S6K1-Myc or S6K2-Myc. Both S6K1 and S6K2 phosphorylated all three eIF5A isoforms, but not the negative control protein CyTPI (cytosolic triose phosphate isomerase), an unrelated protein (Figure 2F). To identify the S6K-targeted phosphorylation site on eIF5A-2, we substituted each of the nine Ser/Thr residues in eIF5A-2 (Ser2, Ser11, Thr18, Thr66, Ser76/Ser77, Thr105/Ser107, and Thr118) with alanine. Subsequent kinase assays using these alanine-substituted variants revealed that Ser2 is specifically phosphorylated by S6K1 (Figure 2G). Collectively, these findings demonstrate that eIF5A is a direct substrate of S6K, phosphorylated at Ser2.

Given that S6Ks phosphorylate *A. thaliana* eIF5A-2 at the Ser2 residue, we next examined whether this N-terminal serine is evolutionarily conserved across major eukaryotic and archaeal lineages. We retrieved eIF5A homologs and their isoforms from Archaea, Fungi, Animals, Protists, and Archaeplastida to assess residue conservation within the N-terminal region. The serine residue corresponding to Ser2 of *A. thaliana* eIF5A-2 is well conserved in Fungi, Protists, and Archaeplastida, whereas this feature is largely absent in Archaea and Animals (Figure S4A). Extending this analysis across Archaeplastida, individual species showed universal conservation of the Ser2-equivalent residue, while eIF5A isoform number varied across clades, generally increasing in land plants relative to green algae (Figure S4B). Multiple sequence alignment of eIF5A homologs and their isoforms from algae and land plants further confirmed that the Ser2-equivalent residue is invariant across the analyzed dataset (119 species, 458 proteins; Figures S4A and S4B). These results indicate that the N-terminal Ser2 residue of eIF5A is under strong evolutionary constraint across the algae-to-land plant lineage, providing an evolutionary basis for S6K-dependent regulation of plant eIF5A.

### eIF5A is required for root growth and root hair formation

To investigate the developmental role of eIF5A, we attempted to disrupt all three *eIF5A* genes simultaneously using a CRISPR-Cas12a construct containing four crRNAs targeting *eIF5A-1*, *eIF5A-2*, and *eIF5A-3* (Figures S5A and S5B). Among 20 screened T1 plants, editing was detected in 4 plants for eIF5A-1 (20 %), 6 plants for eIF5A-2 (30 %), and 9 plants for eIF5A-3 (45 %) (Figure S5C). Bulk Sanger sequencing and Inference of CRISPR Edits (ICE) analysis provisionally classified most edited plants as chimeric or mosaic, with a smaller number classified as putative heterozygotes(Conant et al., 2022). Sanger sequencing chromatograms from two representative T1 plants further showed simultaneous editing at all three eIF5A loci (Figure S5D). Nevertheless, no stable triple knockout line was recovered. These observations suggest that simultaneous disruption of all three eIF5A isoforms severely compromises plant viability or reproductive development. A previous study reported that the *fbr12/eIF5A-2* mutant exhibits extreme dwarfism, reduced organ size and number, and reproductive defects(Feng et al., 2007). Simultaneous loss of all three isoforms may therefore impose a substantially greater developmental burden, consistent with the chimeric and reproductive phenotypes observed in our CRISPR-edited lines.

To overcome this limitation, we generated estradiol-inducible artificial microRNA lines targeting all three *eIF5A* isoforms (*eif5a-ami-es #4* and *eif5a-ami-es #10*) (Figure 3B). Representative images showed that Es-treated *eif5a-ami-es #4* and *eif5a-ami-es #10* seedlings exhibited overall growth defects compared with the corresponding EtOH-treated seedlings (Figure 3A). We first validated *eIF5A* knockdown in these lines. qRT-PCR analysis showed that Es treatment reduced the transcript levels of *eIF5A-1*, *eIF5A-2*, and *eIF5A-3*, and immunoblotting with the anti-eIF5A antibody confirmed a corresponding reduction in eIF5A protein abundance (Figures 3C, 3D, and S5E). These results establish *eif5a-ami-es #4* and *eif5a-ami-es #10* as inducible eIF5A-depletion lines. Estradiol-induced eIF5A depletion caused marked defects in seedling root growth (Figure 3E). We next quantified primary root length and root hair coverage to compare these phenotypes with those observed in TOR- and S6K-perturbed lines. Both *eif5a-ami-es #4* and *eif5a-ami-es #10* showed significantly reduced primary root length and root hair coverage upon Es treatment compared with the EtOH control (Figure 3F). These phenotypes closely resembled those observed in *tor-es* and *s6k1/2-es* lines (Figure 1), indicating that eIF5A is required for the root developmental outputs associated with TOR-S6K signaling.

**Figure 3.**
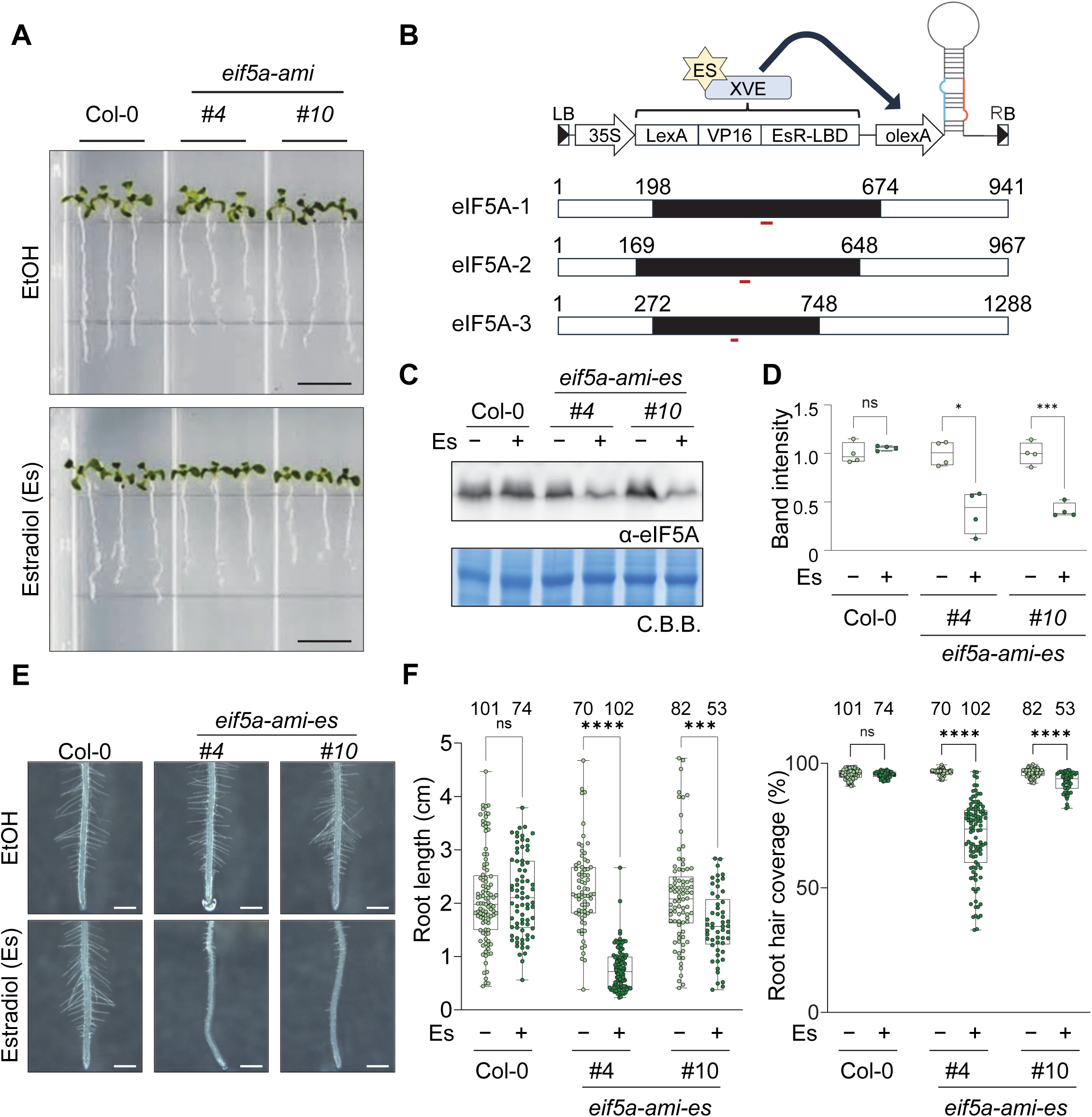
eIF5A depletion impairs primary root growth and root hair coverage. (A) Representative images of Col-0, *eif5a-ami-es #4*, and *eif5a-ami-es #10* seedlings treated with EtOH or 10 μM Es. Scale bars, 1 cm. (B) Schematic illustration of the XVE-based estradiol-inducible artificial microRNA system used to silence *eIF5A-1*, *eIF5A-2*, and *eIF5A-3*. Upon estradiol treatment, activated XVE binds the LexA operator (olexA) and induces *amiRNA* expression. Red bars indicate the amiRNA target sites within each *eIF5A* coding sequence. Numbers indicate nucleotide positions from the transcription start site. (C) Immunoblotting analysis of total eIF5A protein abundance in 7-day-old Col-0, *eif5a-ami-es #4*, and *eif5a-ami-es #10* seedlings treated with EtOH or 10 μM Es. Coomassie Brilliant Blue (CBB) staining shows Rubisco large subunit (rbcL) as a loading control. This experiment was repeated four times with similar results. (D) Quantification of eIF5A band intensity from four independent biological replicates (*n* = 4), one representative result is shown in (C). Band intensities were normalized to the corresponding rbcL band intensity and then expressed relative to the corresponding EtOH control. Boxes indicate the first and third quartiles, the center lines indicate the median, and whiskers indicate the minimum and maximum data points. Individual data points are shown as dots. Statistical significance was assessed using multiple Welch-corrected unpaired t-tests with Holm–Šídák correction for multiple comparisons to compare EtOH and Es within each genotype; ns, P > 0.05; *, P < 0.05; ***, P < 0.001. (E) Representative root images of Col-0, *eif5a-ami-es #4*, and *eif5a-ami-es #10* seedlings under EtOH or 10 μM Es. Scale bars, 1 mm. (F) Box plots represent quantification of primary root length (left) and root hair coverage (right) in Col-0, *eif5a-ami-es #4*, and *eif5a-ami-es #10* seedlings under EtOH or 10 μM Es, corresponding to the roots shown in (E). Boxes indicate the first and third quartiles, the center lines indicate the median, and whiskers indicate the minimum and maximum data points. Sample sizes (*n*) for each group are indicated above each plot. Individual data points are shown as dots. Statistical significance was assessed using multiple Welch-corrected unpaired t-tests with Holm–Šídák correction for multiple comparisons to compare EtOH and Es within each genotype; ns, P > 0.05; ***, P < 0.001; ****, P < 0.0001.

### Developmental functions of *A. thaliana* eIF5A are only partially dependent on hypusination

Hypusination is a unique eIF5A-specific post-translational modification generated through the sequential activities of deoxyhypusine synthase (DHS) and deoxyhypusine hydroxylase (DOHH)(Belda-Palazon et al., 2014; Palfi et al., 2021; Park et al., 2006; Park & Wolff, 2018). In yeast, hypusinated eIF5A promotes translation elongation, including at polyproline motifs(Gutierrez et al., 2013; Saini et al., 2009; Schuller et al., 2017). In mammals, eIF5A-1 and DHS are essential for mouse embryonic development, and hypusinated eIF5A supports translation of Pro-Gly-rich collagen sequences(Barba-Aliaga et al., 2021; Nishimura et al., 2012). We therefore asked whether eIF5A hypusination is similarly required for eIF5A-dependent developmental outputs in *A. thaliana*. We first assessed publicly available *DHS* T-DNA insertion lines, in which T-DNAs are inserted within the 5’ UTR of *DHS* (Figure S6A). Although genotyping PCR confirmed homozygous T-DNA insertions in candidate lines (Figures S6A and S6B), qRT-PCR and immunoblot analyses showed that *DHS* transcript levels, total eIF5A, and hypusinated eIF5A were not substantially reduced in these lines (Figures S6C and S6D), indicating that these T-DNA insertion lines were not suitable for determining the developmental consequences of *DHS* loss of function.

We next compared direct *eIF5A* silencing with *DHS* silencing using virus-induced gene silencing (VIGS). TRV2-eIF5A plants showed pronounced developmental defects compared with TRV2-Myc control plants, including growth retardation and curled leaves (Figures S7A and S7B). TRV2-eIF5A plants also showed reproductive abnormalities, including shortened stamens and petals and thickened carpels (Figure S7C). These reproductive defects were consistent with the seedless-like phenotypes observed in *eIF5A* triple mutant attempts (Figures S5A-S5D). qRT-PCR and immunoblot analyses confirmed reduced levels of transcripts encoding *eIF5A* isoforms and reduced eIF5A protein abundance in TRV2-eIF5A plants (Figures S7D and S7E). These results indicate that eIF5A is required for multiple developmental processes beyond primary root growth. We then silenced *DHS* to test whether perturbing hypusination results in similar phenotype to *eIF5A* depletion. *DHS* silencing caused comparatively mild vegetative phenotypes relative to TRV2-eIF5A plants (Figure S7F). qRT-PCR analysis confirmed reduced expression of the corresponding *DHS* transcripts in TRV2-DHS (Figure S7G). Immunoblot analysis showed that total eIF5A protein levels were comparable between TRV2-Myc and TRV2-DHS plants, whereas hypusinated eIF5A was substantially reduced in TRV2-DHS plants (Figure S7H). Thus, although *DHS* silencing substantially depleted hypusinated eIF5A, it did not recapitulate the developmental defects caused by direct eIF5A depletion. This dissociation suggests that the developmental functions of eIF5A are not uniformly dependent on hypusination, with hypusination requirements likely varying across developmental contexts or stages, consistent with stage-specific requirements for hypusination reported in earlier studies.

We placed this comparison in structural context by examining eIF5A fold conservation and the position of the hypusination loop. The AlphaFold models of *A. thaliana* eIF5A-1 and eIF5A-3 superimposed on the eIF5A-2 crystal structure with Cα RMSD values of 1.796 and 1.584 Å, respectively (Figure S8A)(Teng et al., 2009; Varadi et al., 2024). Yeast eIF5A (PDB 3ER0) and human eIF5A1(Wator et al., 2023) also superimposed on *A. thaliana* eIF5A-2 with Cα RMSD values of 1.680 and 1.938 Å, respectively. Finally, *A. thaliana* eIF5A-2 was superimposed on ribosome-bound hypusinated yeast eIF5A(Schmidt et al., 2016), placing the plant Lys51 side chain in the P-site tRNA CCA-end region in the unrefined model. In the yeast complex, the hypusine tip approached the CCA-end phosphate backbone of the P-site tRNA, whereas the unmodified *A. thaliana* Lys51 side chain was positioned near the CCA-end after superposition (Figure S8C). Although this comparison is based on structural superposition rather than a plant ribosome-bound eIF5A structure, it suggests that *A. thaliana* eIF5A-2 may be geometrically compatible with CCA-end proximity even without hypusine extension. Together, these results show that *A. thaliana* eIF5A is hypusinated, but partial disruption of DHS-dependent hypusination causes milder developmental effects than direct eIF5A depletion, suggesting that the developmental functions of eIF5A are not uniformly dependent on hypusination.

### eIF5A selectively promotes polyproline translation without broadly affecting global translation

eIF5A promotes translation elongation in yeast and in mammalian sequence-specific contexts(Coni et al., 2020; Gutierrez et al., 2013; Saini et al., 2009; Schuller et al., 2017), whereas its contribution to bulk translation in plants has not been directly defined. To address this, we performed a ^35^S-methionine incorporation assay in *eif5a-ami-es* plants (Figures 4A, 4B). *eIF5A* knockdown induced by Es treatment showed no significant difference in ^35^S-methionine incorporation compared to the EtOH-treated control or wild-type plants, indicating that eIF5A does not significantly affect nascent protein synthesis. Polysome profiling similarly revealed no significant alteration in ribosome distribution upon estradiol-induced *eIF5A* knockdown compared to EtOH-treated controls. The comparable monosome and polysome peak profiles suggest that eIF5A depletion does not broadly impair translation initiation, ribosome loading, or elongation at the global level (Figures 4C, 4D). Together, these results indicate that eIF5A is dispensable for global translation in *A. thaliana*.

**Figure 4.**
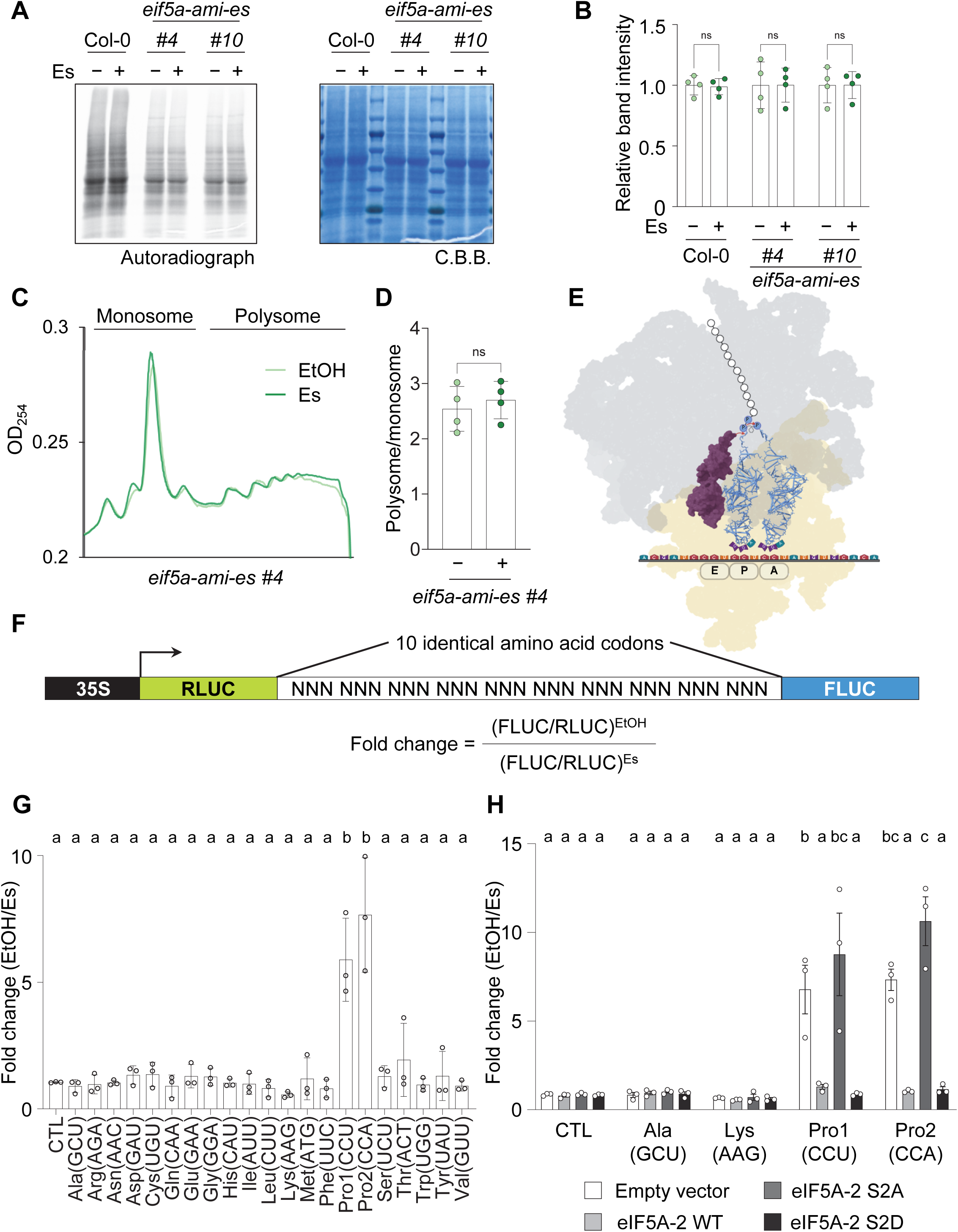
eIF5A selectively promotes polyproline translation through Ser2-dependent activity. (A) ^35^S-methionine incorporation assay in Col-0, *eif5a-ami-es #4*, and *eif5a-ami-es #10* seedlings treated with EtOH or 10 μM Es. Twelve-day-old seedlings were labeled with ^35^S-methionine for 2.5 h under the same treatment conditions. Autoradiography shows newly synthesized proteins, and CBB staining shows protein loading. (B) Quantification of ^35^S-methionine incorporation in (A). Autoradiographic signals were normalized to the corresponding CBB loading signal and are shown relative to the EtOH control. Bar graphs show means ± SD with all data points. *n* = 4 biological replicates. Statistical significance was assessed using multiple Welch-corrected unpaired t-tests with Holm–Šídák correction for multiple comparisons to compare EtOH and Es within each genotype; ns, P > 0.05. (C) Polysome profiling of *eif5a-ami-es #4* seedlings treated with EtOH or 10 μM estradiol. Eight-day-old seedlings were treated with cycloheximide before harvest, and total extracts were separated by sucrose density gradient sedimentation. Absorbance at 254 nm was monitored to visualize monosome and polysome fractions. (D) Quantification of polysome-to-monosome ratios in (C). Bar graphs show means ± SD with all data points. *n* = 4 biological replicates. Statistical significance was assessed using a two-tailed Welch’s t test; ns, P > 0.05. (E) Model for eIF5A function during translation elongation. eIF5A associates near the ribosomal E site and facilitates synthesis of nascent polypeptides containing polyproline stretches. (F) Schematic of the dual-luciferase reporter used to monitor translation through repetitive codon sequences. The reporter contains the 35S promoter, Renilla luciferase (RLUC), 10 identical amino acid codons, and Firefly luciferase (FLUC). Fold change was calculated as (FLUC/RLUC)EtOH divided by (FLUC/RLUC)Es. (G) Dual-luciferase reporter screen in *eif5a-ami-es #4* protoplasts. Reporters containing 10 repeats of the indicated codon were transfected into protoplasts treated with EtOH or 10 μM Es. Fold changes in FLUC/RLUC ratios were normalized to the no-insert control (CTL). Bar graphs show means ± SD with all data points. *n* = 3 biological replicates. Statistical significance was assessed using ordinary one-way ANOVA followed by Tukey’s multiple-comparisons test. Letters indicate statistical groups; groups sharing a letter are not significantly different at P < 0.05. (H) Rescue assay using amiRNA-resistant eIF5A-2 variants. Reporters containing CTL, Ala(GCU), Lys(AAG), Pro1(CCU), or Pro2(CCA) repeats were co-transfected with empty vector, eIF5A-2 WT, eIF5A-2 S2A, or eIF5A-2 S2D in *eif5a-ami-es #4* protoplasts. Fold changes in FLUC/RLUC ratios were normalized as in G. eIF5A-2 WT and S2D restored polyproline reporter output, whereas S2A failed to rescue this defect. Bar graphs show means ± SD with all data points. *n* = 3 biological replicates. Statistical significance was assessed using ordinary one-way ANOVA followed by Tukey’s multiple-comparisons test. Letters indicate statistical groups; groups sharing a letter are not significantly different at P < 0.05.

Since eIF5A and its bacterial functional analog EF-P promote translation in polyproline contexts, with strong stalling at particular XPPX/PPP motifs and substantial sequence-context effects, we tested whether *A. thaliana* eIF5A supports output from a decaproline reporter(Doerfel et al., 2013; Gutierrez et al., 2013; Huter et al., 2017; Peil et al., 2013; Starosta et al., 2014; Ude et al., 2013) (Figure 4E). To examine whether *A. thaliana* eIF5A facilitates translation elongation through repetitive sequence contexts, we employed a dual-luciferase reporter in which Renilla Luciferase (RLUC) and Firefly Luciferase (FLUC) are linked by a stretch of 10 consecutive identical codons under the control of a 35S promoter (Figure 4F). Protoplasts isolated from *eif5a-ami-es #4* plants were transfected with reporters containing a representative codon for each of the 20 amino acids in the repeat region, and treated with Es or EtOH as a control (Figure 4G). Among all amino acids tested, eIF5A knockdown selectively reduced the FLUC/RLUC ratio in the proline repeat constructs (Figure 4G). This reduction was consistently observed with two distinct proline codons (CCU and CCA), suggesting that the effect arises from inefficient translation through repetitive proline stretches rather than codon-specific effects. This finding was further corroborated using TRV2-eIF5A plants, in which the FLUC/RLUC ratio was similarly reduced in the 10-proline repeat construct (Figure S9), supporting the idea that eIF5A facilitates efficient translation of repetitive proline stretches. Collectively, these results indicate that the conserved role of eIF5A in promoting translation elongation through polyproline sequences extends to plants.

To examine whether S6K-mediated phosphorylation of eIF5A at Ser2 is required for its function in polyproline translation, amiRNA-resistant *eIF5A-2-Myc* constructs (WT, S2A, and S2D) carrying silent mutations in the amiRNA target sequence to evade knockdown, were co-expressed with polyproline, polyalanine, or polylysine reporter in *eif5a-ami-es #4* protoplasts. Co-expression of *eIF5A-2 WT-Myc* or the phosphomimetic *S2D* variant selectively restored the FLUC/RLUC ratio in the polyproline reporter, but not in the polyalanine or polylysine controls, whereas the phosphorylation-deficient *S2A* variant failed to rescue translation under any condition (Figures 4H and S10). Collectively, these results demonstrate that eIF5A is required for efficient translation through polyproline sequences and that its phosphorylation status critically modulates this activity.

### Proteomic profiling identifies eIF5A-responsive proteins that support root development

Having established that *A. thaliana* eIF5A promotes translation through polyproline tracts, we asked whether *eIF5A* depletion alters the endogenous proteome. Quantitative proteomic profiling was performed using *eif5a-ami-es #4* seedlings treated with EtOH or Es. Principal component analysis separated the two conditions (Figure S11A). At FDR < 0.01 and absolute log_2_ fold change > 1, 60 proteins decreased and 66 increased after *eIF5A* depletion. The 60 downregulated proteins showed a consistent condition-dependent pattern across the three independently prepared samples per condition (Figure 5A), indicating that partial *eIF5A* depletion altered a defined subset of the proteome rather than uniformly suppressing protein abundance.

**Figure 5.**
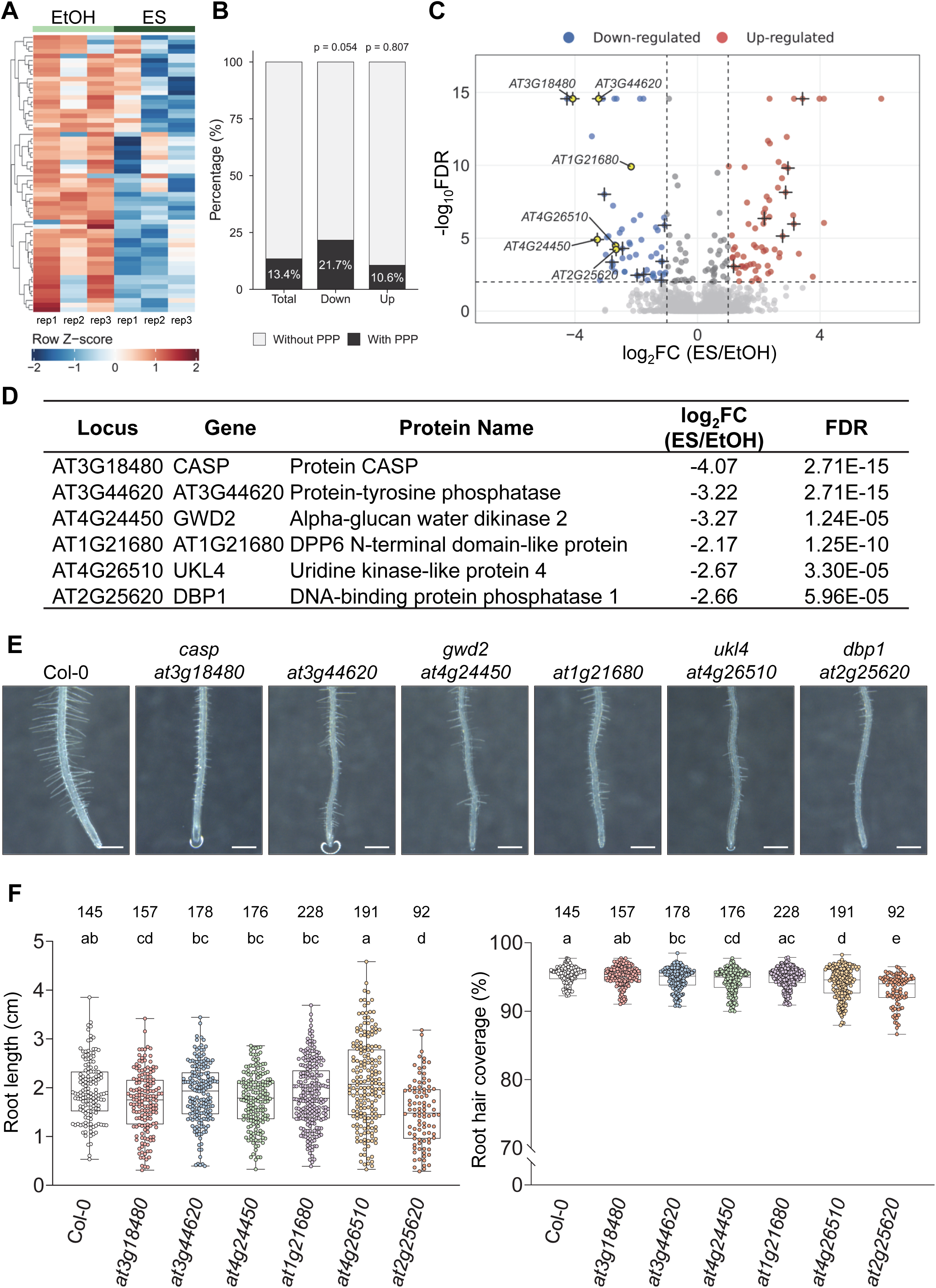
Proteomic profiling identifies eIF5A-responsive proteins that support root development. (A) Heatmap of the 60 proteins with decreased in Es-treated *eif5a-ami-es #4* seedlings relative to EtOH-treated controls (FDR < 0.01 and log_2_ fold change < -1). Values are row-wise Z scores calculated from log_2_-normalized protein abundances from three independently prepared samples per condition. Rows were hierarchically clustered; columns are shown without clustering. (B) Percentages of proteins containing at least one PPP motif among background proteins (the group labeled Total in the current panel; *n* = 4,435), downregulated proteins (Down; *n* = 60), and upregulated proteins (Up; *n* = 66). P values were calculated using one-sided Fisher’s exact tests comparing each regulated set with all other sequence-resolved quantified proteins. (C) Volcano plot of sequence-resolved quantified proteins. The x axis shows log_2_ fold change (ES/EtOH), and the y axis shows -log_10_FDR. Blue and red points indicate downregulated and upregulated proteins, respectively; black plus signs indicate significant PPP-containing proteins. Yellow points identify the six candidates selected for genetic analysis. Dashed lines indicate FDR = 0.01 and |log_2_ fold change| = 1. (D) Locus identifiers, gene names, protein annotations, log_2_ fold changes, and FDR values for the six candidates selected for genetic analysis. (E) Representative images of the primary root differentiation zones of 7-day-old Col-0 and the indicated homozygous insertion mutants. Scale bars, 1 mm. (F) Box plots represent quantification of primary root length (left) and root hair coverage (right) in Col-0, *casp/at3g18480*, *at3g44620*, *gwd2/at4g24450*, *at1g21680*, *ukl4/at4g26510*, and *dbp1/at2g25620*, corresponding to the roots shown in (E). Boxes indicate the first and third quartiles, the center lines indicate medians, and whiskers indicate the minimum and maximum data points. Sample sizes (*n*) for each group are indicated above each plot. Individual data points are shown as dots. Statistical significance was assessed using ordinary one-way ANOVA followed by Tukey’s multiple comparisons test. Letters indicate statistical groups; groups sharing a letter are not significantly different at P < 0.05.

We next tested whether eIF5A-responsive proteins were preferentially associated with proline-rich sequence contexts. TAIR10 sequences were assigned to 4,561 quantified proteins and screened for 16 predefined proline-containing motifs. PPP was present in 13 of 60 downregulated proteins (21.7 %) and 596 of 4,435 background proteins (13.4 %) (Figures 5B and 5C). Although neither motif reached formal statistical significance, PPP- and PGA-containing proteins showed notable numerical enrichment among downregulated proteins. For PPP, this enrichment approached the conventional significance threshold in both a one-sided Fisher’s exact test (P = 0.0537) and a protein-length-adjusted logistic regression (P = 0.0502), although no motif remained significant after multiple-testing correction (Figures S11B-S11D). By contrast, PPP-containing proteins were not enriched among upregulated proteins (7 of 66, 10.6 %; P = 0.8068). Together, the preferential, albeit statistically inconclusive, enrichment of PPP and PGA motifs among downregulated proteins suggests that a subset of proteins containing these proline-rich motifs may be particularly sensitive to *eIF5A* depletion.

To connect these proteomic changes to root development, we examined six strongly downregulated candidates including CASP/AT3G18480, AT3G44620, GWD2/AT4G24450, AT1G21680, UKL4/AT4G26510, and DBP1/AT2G25620 (Figures 5C and 5D). Four candidates contain a PPP motif, whereas AT1G21680 and UKL4 do not. None of the corresponding transcripts decreased after Es treatment in *eif5a-ami-es #4* or *eif5a-ami-es #10*. Instead, *AT3G44620* and *AT4G26510* increased in at least one line (Figure S12). The reduced abundance of these proteins therefore was not explained by lower transcript abundance and is consistent with post-transcriptional regulation.

qRT-PCR confirmed reduced target transcript abundance in each homozygous insertion line (Figure S13). Phenotypic analysis revealed distinct contributions to primary root growth and root hair coverage. *casp/at3g18480* and *dbp1/at2g25620* seedlings had shorter primary roots than Col-0, whereas *at3g44620*, *gwd2/at4g24450*, *ukl4/at4g26510*, and *dbp1/at2g25620* showed reduced root hair coverage (Figures 5E and 5F). *at1g21680* did not show a significant defect in either trait. Thus, eIF5A appears to preferentially support root development through PPP-containing target proteins, although non-PPP targets also contribute to this process.

## Discussion

This study identifies Ser2 as a conserved regulatory site in eIF5A and connects eIF5A to S6K activity, selective translation, and root development in *A. thaliana*. Conditional suppression of TOR, S6K1/2, or the three eIF5A isoforms produced overlapping reductions in primary root length and root hair coverage. S6K1 and S6K2 associated with all three eIF5A isoforms in plant cells, and recombinant S6Ks phosphorylated each isoform in vitro, with Ser2 emerging as the major S6K1-dependent site in eIF5A-2. Depletion of eIF5A also impaired translation of a synthetic deca-proline reporter, and this defect was rescued by amiRNA-resistant wild-type or S2D eIF5A-2, but not by S2A. Together, these findings position eIF5A as a candidate translational output of plant S6K signaling and indicate that the Ser2 state is important for eIF5A activity in the tested context. Whether glucose/TOR controls endogenous eIF5A Ser2 phosphorylation through S6K in vivo remains to be determined.

Plant TOR coordinates translation through several downstream outputs. Glucose-TOR signaling activates root meristems(Li et al., 2017; Xiong et al., 2013). TOR-S6K1-dependent phosphorylation of eIF3h promotes reinitiation on upstream open reading frame-containing mRNAs(Schepetilnikov et al., 2013), whereas TOR-LARP1 signaling changes the translation of defined transcript classes(Scarpin et al., 2020). Our results suggest that eIF5A may define an additional branch through which nutrient-responsive signaling reaches elongation-sensitive sequences. The association between S6Ks and eIF5As, direct phosphorylation in vitro, and the overlapping root phenotypes provide complementary support for this model. Establishing pathway order will require monitoring endogenous eIF5A-pSer2 during glucose withdrawal and repletion and after perturbation of TOR or S6K1/2. Combined with genetic complementation using comparably expressed wild-type, S2A, and S2D proteins, these analyses should distinguish a linear TOR-S6K-eIF5A axis from convergent regulation of root growth and translation.

The strong conservation of Ser2 across the analyzed Archaeplastida sequences indicates functional constraint at the eIF5A N terminus. In maize, CK2 phosphorylates eIF5A at Ser2, and a phosphomimetic substitution alters its nucleocytoplasmic distribution(Łebska et al., 2010; Lewandowska-Gnatowska et al., 2011). In *A. thaliana* leaves, eIF5A-2 and eIF5A-3 Ser2 phosphorylation is higher in the light, declines during extended night, and is elevated in a snrk1α1/α2 mutant, linking the site to cellular energy status(Nukarinen et al., 2016). Our identification of S6K1 and S6K2 as kinases capable of phosphorylating plant eIF5A in vitro extends this regulatory landscape and raises the possibility that Ser2 integrates multiple kinase inputs. The inability of S2A to restore the deca-proline reporter, together with rescue by S2D, is consistent with a functional requirement for phosphorylation competence or negative charge near the N terminus. Because aspartate only approximates phosphoserine, the S2D result supports a charge-dependent regulatory model rather than defining a constitutively phosphorylated active form. Determining endogenous Ser2 occupancy and the effects of this modification on eIF5A localization and ribosome association should clarify how these kinase inputs are translated into eIF5A activity.

The Ser2 result also raises the broader possibility that eIF5A participates in reversible transitions between active and dormant ribosomes. In Trypanosoma cruzi, phosphorylated eIF5A predominates during exponential growth, whereas extensive dephosphorylation accompanies entry into stationary phase. Overexpression of wild-type eIF5A or S2D, but not S2A, increased proliferation and protein synthesis during exponential growth; upon entry into stationary phase, S2D-expressing cells showed less-pronounced translational arrest and reduced viability, supporting a context-dependent phosphorylation-dephosphorylation switch(Chung et al., 2013). Independent cryo-EM structures of Stm1/SERBP1-bound dormant ribosomes also placed eIF5A on the dormant complex, although this work did not examine Ser2(Du et al., 2024). In vertebrate eggs, Dap1b/Dapl1 and eIF5A form one module of a conserved network that stabilizes dormant ribosomes and represses translation(Leesch et al., 2023). In glucose-depleted Schizosaccharomyces pombe, SNOR contacts the hypusinated loop of eIF5A on dormant ribosomes; after glucose repletion, SNOR deletion abolishes polysome recovery, whereas pharmacological inhibition of eIF5A hypusination impairs recovery(Gluc et al., 2026). These studies place eIF5A at multiple stages of the translational cycle, with its role shaped by modification state and binding partners. In *A. thaliana*, whether Ser2 dephosphorylation promotes ribosome dormancy remains an open question. We propose that nutrient sufficiency favors a phosphorylated eIF5A state that supports active translation, whereas nutrient limitation favors dephosphorylation and translational arrest while partner-bound, hypusinated eIF5A helps maintain restart-competent ribosomes. Tracking pSer2, hypusination, and eIF5A distribution across polysomal, 80S, and dormant ribosome fractions during glucose withdrawal and repletion will provide a direct test of this model.

The translational phenotypes further point to selective rather than uniform effects on protein synthesis. eIF5A promotes peptide-bond formation at polyproline motifs and contributes more broadly to elongation and termination(Gutierrez et al., 2013; Pelechano & Alepuz, 2017; Saini et al., 2009; Schuller et al., 2017). In the present assays, eIF5A depletion did not measurably change bulk ^35^S-methionine incorporation or the polysome-to-monosome ratio, yet it reduced output from the deca-proline reporter. Partial or transient depletion could therefore preserve bulk translation while preferentially exposing sequences with a greater elongation requirement for eIF5A. The proteomic data are compatible with this interpretation: 13 of the 60 reduced proteins contained polyproline motifs (21.7 %), compared with 596 of 4,435 quantified background proteins (13.4 %). However, this enrichment was borderline in the one-sided Fisher test (P = 0.0537), and no tested motif remained significant after multiple-testing correction. It should therefore be viewed as a trend rather than a statistically established motif rule. Ribosome profiling with codon-resolution pausing analysis, ideally paired with nascent-chain proteomics, should define the endogenous target spectrum and distinguish direct elongation defects from secondary changes in protein stability or cellular state.

The candidate-gene analysis provides a bridge between translational control and root development. The six selected proteins decreased without a corresponding reduction in transcript abundance, consistent with post-transcriptional regulation, and five of the six insertion lines altered primary root length, root hair coverage, or both. These results nominate multiple proteins that may transmit eIF5A-dependent translational changes to root growth. The inclusion of UKL4, which lacks the tested polyproline motifs, further suggests that eIF5A-sensitive translation may extend beyond a single sequence class. Mutant phenocopy alone does not assign direct translational dependence, but transcript-specific ribosome occupancy or nascent-protein measurements, followed by restoration of candidate expression in eIF5A-depleted plants, should distinguish direct effectors from genes that influence the same traits in parallel. Editing of implicated stalling motifs, where this can be achieved without altering protein function, would provide an additional test of sequence-dependent regulation.

Among these candidates, DBP1 provides the clearest prior connection to root developmental regulation. DBP1 is a PPP-containing, plant-specific DNA-binding protein phosphatase of the PP2C class(Carrasco et al., 2005; Carrasco et al., 2014; Castello et al., 2010). AtDBP1 is auxin inducible; its expression is enhanced in the root-apex cell-division zone, and its promoter is active in the root meristem, lateral root initiation sites, and vascular tissues; antisense suppression attenuates auxin responsiveness and auxin-induced callus formation(Alliotte et al., 1989; Bernstein et al., 2015). An earlier analysis of the same dbp1 insertion line detected no gross whole-plant growth defect, whereas our quantitative seedling assay revealed both shorter primary roots and reduced root hair coverage, consistent with a subtle, stage- or tissue-specific developmental role(Castello et al., 2010). These observations make DBP1 a biologically grounded candidate through which eIF5A-sensitive translation could influence auxin-associated root growth, while direct translational dependence can now be tested by measuring DBP1 ribosome occupancy or nascent synthesis after eIF5A depletion. Ser2 phosphorylation should be considered alongside hypusination, the defining modification of eIF5A. Hypusinated eIF5A promotes elongation(Saini et al., 2009), and independent ribosome-bound yeast structures place the hypusine side chain next to A76 of the P-site tRNA, consistent with stabilization of the peptidyl-transfer geometry(Melnikov et al., 2016; Schmidt et al., 2016). In *A. thaliana*, eIF5A hypusination is environmentally regulated, including by abscisic acid(Belda-Palazon et al., 2014), and conditional suppression of deoxyhypusine synthase disrupts multiple developmental traits, including root architecture and root hair formation(Belda-Palazon et al., 2016). In our experiments, virus-induced DHS silencing produced a milder phenotype than direct eIF5A silencing but left detectable modified eIF5A. This outcome is most consistent with partial or context-dependent suppression and therefore does not argue against an important contribution of hypusination to root development. The structural superposition further places the unmodified *A. thaliana* Lys51 side chain in a sterically unfavorable position relative to the P-site tRNA. Because the 0.4 Å value derives from an unrefined comparative model, it should be interpreted as geometric incompatibility rather than as a physical contact distance. Nevertheless, the clash supports the conserved structural importance of modifying Lys51, which can ultimately be resolved by a plant eIF5A-ribosome structure. Joint analysis of Ser2 phosphorylation and Lys51 hypusination should reveal whether these modifications act independently or coordinately during active translation and dormancy-restart transitions.

Finally, the three *A. thaliana* eIF5A isoforms are likely to combine shared and specialized functions. All three associated with S6Ks and were phosphorylated in vitro, whereas their simultaneous depletion produced strong root defects. Prior studies link eIF5A-2 to root protoxylem development and cytokinin signaling(Ren et al., 2013), and *A. thaliana* eIF5As to pathogen-induced cell death(Hopkins et al., 2008). The inability to recover a triple-edited line is consistent with an essential shared function, although isoform-specific complementation under native or root-zone-specific promoters will be needed to resolve redundancy from specialization. Coupling these experiments to measurements of Ser2 phosphorylation, hypusination, and ribosome association should further define how individual isoforms respond to nutrient and developmental signals. Overall, our findings identify Ser2 as a conserved S6K-responsive site in vitro and establish the importance of eIF5A-dependent translation for the measured root traits. They support a model in which TOR-S6K signaling, eIF5A modification, and selective translation intersect to coordinate root development, and provide a framework for determining how glucose/TOR controls eIF5A phosphorylation in vivo.

## Material and Methods

### Plant Materials and Growth Conditions

*Arabidopsis thaliana* Columbia-0 (Col-0) was used as the wild-type background. Seeds were surface sterilized with 70 % EtOH three times, rinsed with three times with sterile water, and and stratified at 4 °C for 3 days. Unless otherwise stated, plants were maintained at 22 °C under a 16-h light/8-h dark photoperiod at 120-150 μmol m^-2^ s^-1^ using Philips TLD36W/865/FL40SS/36/EX-D lamps.

For liquid culture, seeds were sown in six-well plates containing 1 mL per well of 0.5X Murashige and Skoog (MS) basal salt medium (Duchefa Biochemie, M0221) adjusted to pH 5.7 with KOH. After germination, seedlings were transferred to the same medium supplemented with 30 mM glucose (Sigma-Aldrich, 84097), and the medium was replaced every 2-3 days. This culture regime was adapted from previous study(Lee et al., 2017).

A 30 mM β-estradiol stock solution (Sigma-Aldrich, E8875) was prepared in 100 % EtOH. β-estradiol was added to the medium at 10 μM, and an equal volume of EtOH was used as the solvent control. Unless indicated otherwise, induction was initiated at 5 days after germination (DAG). The XVE-based inducible system was used as described previously(Zuo et al., 2000).

For standard solid culture, seeds were sown on 0.5X MS medium containing vitamins (Duchefa Biochemie, M0222), 30 mM sucrose (Duchefa Biochemie, S0809), and 0.7 % phyto agar (Duchefa Biochemie, P1003), adjusted to pH 5.7 with KOH.

*Nicotiana benthamiana* plants used for transient protein expression were grown at 22 °C under a 16-h light/8-h dark photoperiod at approximately 100-120 μmol m^-2^ s^-1^.

The estradiol-inducible *TOR* RNAi lines *tor-es #1* and *tor-es #2* were described previously(Xiong & Sheen, 2012). The estradiol-inducible multi-amiRNA lines *s6k1/2-es #2* and *s6k1/2-es #11*, targeting *S6K1* (AT3G08730) and *S6K2* (AT3G08720), and *eif5a-ami-es #4* and *eif5a-ami-es #10*, targeting *eIF5A-1* (AT1G13950), *eIF5A-2* (AT1G26630), and *eIF5A-3* (AT1G69410), were generated in the Col-0 background in this study. A *35S::Flag-S6K1* line was also generated in Col-0.

The DHS insertion lines GK-727A01 and SALK_113035C (AT5G05920), and the candidate insertion lines *casp/at3g18480* (SALK_076562), *at3g44620* (SALK_101423C), *gwd2/at4g24450* (GK-257E09), *at1g21680* (SALKseq_043847.1), *ukl4/at4g26510* (SAIL_895_A10), and *dbp1/at2g25620* (SALK_005240C) were obtained from ABRC.

### Plasmid construction

All coding sequences were amplified from cDNA with Phusion High-Fidelity DNA Polymerase (New England Biolabs, M0530) using the primers listed in Table S1. Two cloning workflows were used. PCR products amplified with long primers containing vector-overlap sequences were inserted directly into linearized destination vectors with the Overlap Cloner DNA Cloning Kit (Elpis Biotech, EBK-1012) according to the manufacturer’s instructions. PCR products amplified with short restriction-site primers were first cloned into pJET1.2 (Thermo Fisher Scientific, K1231), verified by Sanger sequencing, excised with the indicated restriction enzymes, and ligated into the corresponding digested destination vector. All final coding sequences and vector-insert junctions were verified by Sanger sequencing.

For transient expression, Flag-S6K1 and Flag-CDC123 were generated by overlap cloning into pCAMBIA1390-Flag linearized with SalI (Thermo Fisher Scientific, FD0644) and BcuI/SpeI (Thermo Fisher Scientific, FD1254), whereas Flag-eIF5A-1, Flag-eIF5A-2, and Flag-eIF5A-3 were cloned into the same backbone linearized with SalI and EcoRI (Thermo Fisher Scientific, FD0274). S6K1-Myc and S6K2-Myc were generated using pCAMBIA1390-Myc linearized with SalI and BglII (Thermo Fisher Scientific, FD0084)(Ahn et al., 2015). CDC123-VYNE, S6K1-VYNE, S6K2-VYNE, eIF5A-1-VYCE, eIF5A-2-VYCE, and eIF5A-3-VYCE were generated by overlap cloning into the corresponding pVYNE or pVYCE backbone(Waadt et al., 2008) linearized with BamHI (Thermo Fisher Scientific, FD0054) and XhoI (Thermo Fisher Scientific, FD0694). All constructs were assembled using the primers listed in Table S1. The pVYNE and pVYCE vectors were used as described previously(Waadt et al., 2008).

The *eIF5A-1*, *eIF5A-2*, and *eIF5A-3* coding sequences were amplified using primers containing SalI and EcoRI sites, cloned into pJET1.2, excised with SalI and EcoRI, and ligated into pCAMBIA1390-Myc digested with the same enzymes using T4 DNA ligase (Thermo Fisher Scientific, EL0011).

For recombinant protein expression, the coding sequences of eIF5A-1, eIF5A-2, and eIF5A-3 were assembled into pET-29a(+) linearized with BamHI and HindIII to express proteins carrying a C-terminal His tag. The eIF5A-2 S2A and S11A variants were generated using mutagenic full-length forward primers and a common reverse primer. The T18A, T66A, S76A/S77A, T105A/S107A, and T118A variants were generated by overlap-extension site-directed mutagenesis using the listed internal primer pairs and subsequently assembled into BamHI- and HindIII-linearized pET-29a(+), producing recombinant proteins with C-terminal His tags. His-CyTPI served as a negative-control substrate.

For VIGS constructs, a triple Myc epitope sequence was inserted into pTRV2 linearized with EcoRI and XhoI to generate the pTRV2-Myc control construct. The full-length *eIF5A-2* coding sequence was amplified and assembled into pTRV2 linearized with EcoRI and XhoI. For pTRV2-DHS, the fragment corresponding to nucleotides 1-593 of the DHS coding sequence was amplified and assembled into pTRV2 linearized with EcoRI. All constructs were generated using the Elpis Overlap Cloner system.

For dual-luciferase reporters, complementary oligonucleotides encoding ten consecutive identical codons were annealed to generate double-stranded inserts(Gutierrez et al., 2013). p326-RLUC-FLUC was digested with SmaI (Thermo Fisher Scientific, FD0664) to generate blunt ends, and each annealed insert was ligated between RLUC and FLUC using T4 DNA ligase. The two possible insert orientations generated the codon pairs represented by shared oligonucleotide names in Table S1. Reporters insert sequences and orientations were verified by Sanger sequencing. AmiRNA-resistant eIF5A-2-Myc WT, S2A, and S2D constructs were generated by overlap cloning into p326-Myc linearized with XhoI and EcoRI. To confer amiRNA resistance without altering the encoded amino acid sequence, nucleotides 226-246 of the *eIF5A-2* coding sequence were recoded from 5′-TCTTCCCACAATTGTGATGTT-3′ to 5′-AGTAGTCATAACTGCGATGTG-3′.

The *S6K1/2* and *eIF5A* multi-amiRNA sequences were introduced into the MIR319a precursor in pRS300 by overlap PCR as described previously(Schwab et al., 2006). The resulting amiRNA cassettes were inserted into pER8 and pER10, respectively, after linearization with XhoI and BcuI/SpeI, using the pER8_10-ami_F and pER8_10-ami_R primers listed in Table S1 and the Elpis Overlap Cloner system.

The multiplex pAGE2-C12 T-DNA construct contained pNOS::NPTII-OCSt, pRPS5A::ttLbCas12a-EUv1ter, and a U6 promoter-driven array of four crRNAs. The temperature-tolerant LbCas12a variant was described previously(Schindele & Puchta, 2020). The source or construction of pAGE2-C12 and EUv1ter should be stated separately from the ttLbCas12a citation. crRNA1 and crRNA2 targeted *eIF5A-1*, crRNA3 targeted *eIF5A-2*, and crRNA4 targeted *eIF5A-3*. Guide sequences are listed in Table S1.

### Generation of transgenic *A. thaliana* plants

All binary vectors were introduced into *Agrobacterium tumefaciens* C58C1. Agrobacterial cultures were grown overnight at 28 °C in YEP medium (LPS Solution, YEP-05)

supplemented with rifampicin (Duchefa Biochemie, R0146) and kanamycin (Duchefa Biochemie, K0126), each at 50 μg mL^-1^. Cells were harvested and resuspended to an OD600 of 1.0 in infiltration solution containing 5 % (w/v) sucrose and 0.03 % (v/v) Silwet L-77. Col-0 plants were transformed using the floral-dip method(Clough & Bent, 1998; Lee et al., 2017). T1 plants carrying the *35S::Flag-S6K1* or *S6K1/2* multi-amiRNA construct were selected on medium containing 30 μg mL^-1^ hygromycin (Duchefa Biochemie, H0192), whereas those carrying the *eIF5A* multi-amiRNA construct were selected using 35 μg mL^-1^ kanamycin. At least 30 independent T1 transformants were initially screened for each construct, and one or two independent lines were selected for subsequent analyses. One *35S::Flag-S6K1* line, *s6k1/2-es #2* and *s6k1/2-es #11*, and *eif5a-ami-es* #4 and *eif5a-ami-es #10* were used in this study.

### Virus-induced gene silencing

TRV-based VIGS was performed as described previously(Ahn et al., 2019; Burch-Smith et al., 2006; Lee et al., 2017), with the modifications described below. *A. tumefaciens* C58C1 cultures carrying pTRV1, pTRV2-Myc, pTRV2-eIF5A, or pTRV2-DHS were grown overnight at 28 °C in YEP medium containing kanamycin and rifampicin, each at 50 μg mL^-1^. Cells were harvested and resuspended to an OD600 of 1.0 in infiltration buffer containing 10 mM MES-KOH (pH 5.6), 10 mM MgCl_2_, and 0.2 mM acetosyringone. Following resuspension, the cultures were incubated for 4 h. The pTRV1 culture was then mixed with each pTRV2 culture at a 1:1 ratio and infiltrated into *A. thaliana* Col-0 plants at the two-true-leaf stage. Vegetative and reproductive phenotypes were documented at 18, 21, and 30 days after infiltration (DAI), and the ninth leaf was harvested at 15DAI for qRT-PCR and immunoblotting.

### Agroinfiltration for transient protein expression

*A. tumefaciens* C58C1 cultures carrying the indicated expression constructs or P19 were grown overnight at 28 °C in YEP medium containing kanamycin and rifampicin, each at 50 μg mL^-1^(Lee et al., 2017). Cells were harvested and resuspended to an OD600 of 1.0 in infiltration buffer containing 10 mM MES-KOH (pH 5.6), 10 mM MgSO_4_, and 0.5 mM acetosyringone. Following resuspension, the cultures were incubated for 4 h. For expression of a single construct, the corresponding culture was mixed with the P19 culture at a 1:1 ratio. For co-expression of two constructs, the two expression cultures and the P19 culture were mixed at a 1:1:1 ratio. The resulting mixtures were infiltrated into *N. benthamiana* leaves. Leaves were harvested or imaged 3 days after infiltration.

### Primary-root and root-hair phenotyping

Seeds were sown on 0.25X MS medium containing vitamins and 1 % Gelrite (Duchefa Biochemie, G1101), adjusted to pH 5.7 with KOH, and grown vertically for 7 days at 22 °C under continuous light (120-150 μmol m^-2^ s^-1^). Roots were imaged using a stereomicroscope (Leica Microsystems, M50). Primary-root length and root-hair coverage were quantified using ImageJ version 1.54k(Schneider et al., 2012). Root-hair coverage was calculated as the length of the primary root occupied by visible root hairs divided by the total primary-root length and multiplied by 100.

### Flag-S6K1 affinity purification and LC-MS/MS

Col-0 and *35S::Flag-S6K1* seedlings were grown for 7 days under light and glucose (LG) conditions in 0.5X MS medium supplemented with 30 mM glucose. Col-0 seedlings were harvested at 7 days after germination (DAG) and used as the affinity-purification background control. The *35S::Flag-S6K1* seedlings were transferred to darkness and glucose-free medium for 24 h (darkness and starvation, DS) and harvested immediately or after 6 h of recovery under LG conditions. Two independently prepared biological samples were analyzed per condition.

Tissue was homogenized in extraction buffer containing 50 mM Na_3_PO_4_ (pH 7.4), 150 mM NaCl (Duchefa Biochemie, S05820), 10 % glycerol (Duchefa Biochemie, G1345), 5 mM EDTA (Sigma-Aldrich, E5134), 1 mM DTT (Duchefa Biochemie, D1309), 1 % Triton X-100, ReadyShield phosphatase inhibitor cocktail (Sigma-Aldrich, P0001), and cOmplete EDTA-free protease inhibitor cocktail (Roche, 11873580001). Lysates were clarified by centrifugation at 20,000 × g for 15 min at 4 °C, and protein concentrations were determined using the Bradford assay(Bradford, 1976) (Bio-Rad, 5000006). Equal amounts of protein were incubated with 30 μL EZview Red anti-Flag M2 affinity gel (Sigma-Aldrich, F2426) for 4 h at 4 °C. The beads were washed three times with extraction buffer containing 0.1 % Triton X-100.

Bound proteins were eluted by heating the beads in 5× SDS loading buffer for 10 min at 70 °C. Twenty microliters of each eluate was loaded per lane and separated by SDS-PAGE using a 5 % stacking gel and a 10 % resolving gel. Electrophoresis was performed at 60 V through the stacking gel and 100 V through the resolving gel. Gels were stained for 15 min with InstantBlue Coomassie Protein Stain (Abcam, AB119211). The entire stained portion of each of the six sample lanes, encompassing all visible protein bands, was excised and processed separately for LC-MS/MS analysis.

Gel pieces were washed with water and destained four times for 15 min in a 1:1 mixture of methanol and 0.1 M ammonium bicarbonate (ABC, pH 7.8). The gel pieces were washed with 0.1 M ABC/acetonitrile for 5 min, dehydrated twice with acetonitrile, and dried. Proteins were reduced with 25 mM DTT at 56 °C for 20 min and alkylated with 55 mM iodoacetamide for 20 min at room temperature in darkness. The gel pieces were washed, dehydrated, and dried again. Sequencing Grade Modified Trypsin (Promega, V5111) was added to the gel pieces and allowed to absorb for 10 min. Digestion was then performed in 50 mM ABC for 16 h at 37 °C. Peptides were extracted twice with 80 % acetonitrile for 5 min, and the combined extracts were dried completely using a SpeedVac. Peptides were reconstituted in 0.1 % formic acid by vortexing for 30 min and clarified by centrifugation at 13,000 rpm for 10 min. The supernatants were transferred to injection vials for LC-MS/MS analysis.

Peptides were analyzed using an Ultimate 3000 HPLC system coupled to a Q Exactive Plus mass spectrometer (Thermo Fisher Scientific). Samples were loaded onto a trap column (Thermo Fisher Scientific, 164199, 100 μm internal diameter, 150 mm length, 100 Å, 5 μm) and separated on an analytical column (Thermo Fisher Scientific, 164569, 75 μm internal diameter, 250 mm length, 100 Å, 3 μm). Mobile phase A consisted of water containing 0.1% formic acid, and mobile phase B consisted of acetonitrile containing 0.1 % formic acid. Five microliters of each sample was injected, and separation was performed at 0.3 μL min^-1^ and 25 °C. The percentage of mobile phase B was programmed as follows: 5 % from 0 to 5 min, 5-15 % from 5 to 12 min, 15-60 % from 12 to 45 min, 60-95 % from 45 to 47.5 min, 95 % from 47.5 to 52.5 min, 95-5 % from 52.5 to 55 min, and 5 % from 55 to 60 min.

Full-MS and data-dependent HCD MS2 spectra were acquired in positive-ion mode. Full-MS scans were acquired over a range of 150-2,000 m/z at a resolution of 70,000 with an AGC target of 1 × 10^6. MS2 spectra were acquired at a resolution of 35,000 with an AGC target of 5 × 10^4 and a maximum ion injection time of 50 ms. The cycle time was 3 s, the isolation window was 2.0 m/z with a 0.6 m/z offset, the fixed first mass was 100 m/z, and the normalized HCD collision energy was 30 %.

Raw files were reprocessed using MaxQuant v2.4.2.0(Cox & Mann, 2008) with the Andromeda search engine(Cox et al., 2011). Spectra were searched against an *A. thaliana* UniProtKB FASTA database (taxonomy ID 3702) downloaded on July 26, 2023. Reverse decoy sequences and the MaxQuant contaminant database were included in the search. Trypsin/P was specified as the protease, with up to two missed cleavages allowed. The minimum peptide length was seven amino acids, and up to five modifications per peptide were permitted. Carbamidomethylation of cysteine was specified as a fixed modification. Oxidation of methionine and acetylation of the protein N terminus were specified as variable modifications. The precursor mass tolerances were 20 ppm for the first search and 4.5 ppm for the main search. The FTMS MS2 fragment-ion match tolerance was 20 ppm. Peptide, protein, and modification-site false discovery rates were set to 1 %. At least one peptide, including at least one razor peptide, was required for protein identification. Match between runs and label-free quantification were disabled.

Within each condition, proteins detected in both independently prepared biological samples were retained. The protein list obtained from the two Col-0 samples was used as the affinity-purification background and subtracted from the corresponding S6K1-DS and S6K1-LG protein lists. Proteins remaining in at least one *35S::Flag-S6K1* condition were considered candidate S6K1-associated proteins.

### Generation of Anti-eIF5A-2 Polyclonal Antibody

A rabbit polyclonal antibody against *A. thaliana* eIF5A-2 was generated as a custom service (AbFrontier) using two New Zealand white rabbits immunized with recombinant His-eIF5A-2 antigen (total 4 mg). Serum from the second boost was used without further purification.

### Protein extraction and immunoblotting

Total protein was extracted from *A. thaliana* seedlings or leaves and *N. benthamiana* leaves using the Na_3_PO_4_-based extraction buffer described above. Extracts were clarified by centrifugation at 20,000 × g for 15 min at 4 °C, and protein concentrations were determined using the Bradford assay (Bio-Rad, 5000006). Equal amounts of protein were mixed with SDS sample buffer and separated by SDS-PAGE. Proteins were wet-transferred to PVDF membranes (Millipore, ISEQ85R) at 90 V for 70 min using 10 mM CAPS (Sigma-Aldrich, C2632) transfer buffer adjusted to pH 11 with NaOH.

Membranes used for eIF5A or hypusine detection were blocked for 30 min at room temperature in TTBS containing 5 % (w/v) BSA (LPS Solution, BSA100). The corresponding primary and secondary antibodies were diluted in the TTBS. Membranes used for all other targets were blocked for 30 min at room temperature in TTBS containing 5 % (w/v) skim milk (LPS Solution, SKI500), which was also used for antibody dilution.

Primary antibodies were anti-eIF5A (1:1,000, generated in this study), anti-hypusine (1:1,000, ProteoGenix, PTX18841), anti-GFP (1:5,000, Roche, 11814460001), Flag-HRP (1:5,000, Sigma-Aldrich, A8592), Myc-HRP (1:5,000, Upstate, 16-213), and HA-HRP (1:5,000, Roche, 12013819001). HRP-conjugated anti-rabbit (1:5,000, Upstate, 12-348) or anti-mouse (1:5,000, Upstate, 12-349) secondary antibodies were used for unconjugated primary antibodies. Signals were developed using Western Pico ECL Kit (LPS Solution, PICO-250) and detected with an ImageQuant LAS 4000 system (GE Healthcare Life Sciences). Band intensities were quantified using ImageJ. Relative hypusination was calculated by dividing the anti-hypusine signal by the total eIF5A signal. Where indicated, total eIF5A was normalized to the corresponding Coomassie-stained Rubisco large-subunit band.

### Co-immunoprecipitation (Co-IP)

Flag-S6K1 or Flag-CDC123 was co-expressed with eIF5A-1-Myc, eIF5A-2-Myc, or eIF5A-3-Myc in *N. benthamiana* leaves(Ahn et al., 2015; Lee et al., 2017). Flag-CDC123 served as a negative control. At 3 days after infiltration, proteins were extracted using the Na_3_PO_4_-based extraction buffer described above. Equal amounts of total protein were incubated with 30 μL EZview Red anti-Flag M2 affinity gel for 4 h at 4 °C. The beads were washed three times with extraction buffer containing 0.1 % Triton X-100. Bound proteins were eluted by boiling in SDS sample buffer and analyzed by immunoblotting with anti-Flag and anti-Myc antibodies.

For semi-native co-immunoprecipitation, Col-0 and *35S::Flag-S6K1* seedlings were grown for 5 days after germination on glucose-containing MS medium (MSG). For the glucose-supplied condition, seedlings were subsequently grown for an additional 5 days on MSG supplemented with Es or the corresponding EtOH. For the glucose-depleted condition, seedlings were grown for 4 days on MSG supplemented with Es and then transferred to glucose free MS medium containing ES and maintained in darkness for 24 h. Protein extracts were incubated with anti-eIF5A serum at a 1:100 dilution and 20 μL Protein A agarose (Thermo Fisher Scientific, 20333). The beads were washed three times, and bound proteins were eluted by boiling in SDS sample buffer for 5 min before immunoblotting with anti-eIF5A and anti-Flag antibodies. This assay was used to assess co-precipitation but not to quantify glucose-dependent changes in interaction strength because the panel did not include input-normalized IP quantification.

### Bimolecular fluorescence complementation

Bimolecular fluorescence complementation (BiFC) was performed with pVYNE/pVYCE vectors as described previously(Ahn et al., 2015; Waadt et al., 2008), with the modifications described here. CDC123-VYNE, S6K1-VYNE, or S6K2-VYNE was co-expressed with eIF5A-1-VYCE, eIF5A-2-VYCE, or eIF5A-3-VYCE in leaves of 3-week-old *N. benthamiana*. CDC123-VYNE served as a negative control. Venus/YFP fluorescence was examined 3 days after infiltration with a Zeiss LSM 700 confocal laser-scanning microscope. Images compared within an experiment were acquired with identical settings.

### Recombinant protein expression and purification

pET-29a(+) constructs encoding wild-type or mutant eIF5A proteins were introduced into *Escherichia coli* BL21. Cultures were grown at 37 °C to an OD600 of 0.4, shifted to 16 °C, and induced with 0.25 mM IPTG for 16 h. Cells were harvested by centrifugation at 4,000 × g for 15 min at 4 °C. Cell pellets were resuspended in purification buffer containing 20 mM Tris-Cl (pH 7.5), 200 mM NaCl, and 30 mM imidazole. Cells were disrupted by sonication at 40 % amplitude using 7-s pulses separated by 7-s intervals for a total sonication-on time of 10 min. Lysates were clarified by centrifugation at 15,000 × g for 30 min at 4 °C.

Cleared lysates were applied to His60 Ni Superflow resin (Clontech, 635659). The resin was washed with purification buffer containing 20 mM Tris-Cl (pH 7.5), 200 mM NaCl, and 30 mM imidazole. Bound proteins were eluted using buffer containing 20 mM Tris-Cl (pH 7.5), 200 mM NaCl, and 300 mM imidazole. Eluted proteins were concentrated using Amicon Ultra-0.5 centrifugal filter units with a 10-kDa molecular-weight cutoff and regenerated cellulose membrane (Millipore, UFC5010). The buffer was exchanged into 1× PBS through three successive rounds of PBS addition and reconcentration.

Protein concentrations were determined using the Bradford assay (Bio-Rad, 5000006). Protein purity was assessed by 10 % SDS-PAGE followed by staining with Sun-Gel Staining Solution (LPS Solution, SGS01). Purified proteins were stored in 1× PBS at approximately 3 mg mL^-1^ at -80 °C. The same purification and buffer-exchange procedure was used for all wild-type and mutant eIF5A proteins analyzed in Figures 2F and 2G.

### In vitro kinase assay

In vitro kinase assays were performed as described previously(Lee et al., 2023; Lee et al., 2017), with modifications. S6K1-Myc and S6K2-Myc were transiently expressed in *N benthamiana*. Equal amounts of extract were incubated for 4 h at 4 °C with EZview Red anti-c-Myc affinity gel (Sigma-Aldrich, E6654) and washed three times. Kinase reactions (20 μL) contained immunoprecipitated S6K1-Myc or S6K2-Myc, 10 μg recombinant substrate, 10 μCi [γ-32P]ATP (PerkinElmer, Waltham, MA, USA), 10 μM nonradioactive ATP, 20 mM HEPES (Cytiva, SH30237), 125 mM NaCl, 10 mM MgCl_2_ (Duchefa Biochemie, M0533), and 5 mM MnCl_2_ (Sigma-Aldrich, M5005). Reactions were incubated for 30 min at 30 °C, stopped with 2× SDS sample buffer, and heated for 10 min at 70 °C.

Proteins were separated by SDS-PAGE, stained with Sun-Gel staining solution (LPS Solution, SGS01), dried, and analyzed by phosphorimaging with a BAS-2500 bio-imaging analyzer (Fujifilm). Kinase loading was assessed by anti-Myc-HRP immunoblotting. His-CyTPI served as a negative-control substrate.

### RNA extraction and qRT-PCR

Total RNA was isolated from the indicated seedlings or VIGS leaves using the IQeasy Plus Plant RNA Extraction Mini Kit (iNtRON Biotechnology, 17491). First-strand cDNA was synthesized from 1 μg total RNA using the RevertAid First Strand cDNA Synthesis Kit (Thermo Fisher Scientific, K1622) with the oligo(dT)18 primer supplied with the kit according to the manufacturer’s instructions. qRT-PCR was performed using PowerUp SYBR Green Master Mix (Applied Biosystems, A25741) on a StepOnePlus Real-Time PCR System (Applied Biosystems). Reaction setup, amplification, and melt-curve analysis were performed according to the manufacturer’s instructions. Relative transcript abundance was calculated using the 2^-ΔΔCt^ method(Livak & Schmittgen, 2001) and normalized to *PP2AA3*. Primer sequences are listed in Table S1.

### ^35^S-methionine incorporation

^35^S-methionine incorporation was measured as described previously(Ahn et al., 2011; Lee et al., 2017), with modifications. Nine-day-old liquid-grown seedlings were transferred to medium containing EtOH or 10 μM β-estradiol for 3 days and then labeled for 2.5 h under the same treatment condition with 50 μCi ^35^S-methionine (PerkinElmer, Waltham, MA, USA). Seedlings were washed twice with culture medium, and total protein was extracted. Protein concentration was determined by Bradford assay, and equal protein amounts were loaded on 4 %-20 % gradient SDS-PAGE gels. Gels were stained with Sun-Gel, dried, and analyzed with a BAS-2500. Radioactive and Coomassie band intensities from the same 35-75 kDa region were measured in ImageJ, and incorporation was calculated as the radioactive signal divided by the corresponding Coomassie signal. Four independent biological replicates were analyzed.

### Polysome profiling

Polysome profiling was performed as described previously (Lee et al., 2017), with modifications. Five-day-old liquid-grown seedlings were treated with EtOH or 10 μM β-estradiol for 3 days and incubated with 50 μg mL^-1^ cycloheximide for 5 min immediately before harvest. Frozen tissue (0.2 g) was homogenized in 1 mL polysome-isolation buffer containing 200 mM Tris-HCl (pH 8.4), 50 mM KCl, 25 mM MgCl_2_, 1 % sodium deoxycholate (Sigma-Aldrich, D6750), 2 % polyoxyethylene (10) tridecyl ether (Sigma-Aldrich, P2393), 400 U mL^-1^ RiboLock RNase inhibitor (Thermo Fisher Scientific, EO0381), and 50 μg mL^-1^ cycloheximide. Extracts were clarified by centrifugation at 20,000 × g for 15 min at 4 °C. Five hundred microliters of each supernatant was layered onto an 11.5 mL linear 15-50 % (w/v) sucrose gradient prepared in buffer containing 200 mM Tris-HCl (pH 8.4), 50 mM KCl, and 25 mM MgCl_2_.

Gradients were centrifuged at 38,000 rpm for 3.5 h at 4 °C in a Beckman SW41 Ti rotor. Absorbance at 254 nm was recorded using a BioLogic low-pressure chromatography system (Bio-Rad, 7318305EDU). Monosome and polysome peak areas were integrated using ImageJ. Four independent biological replicates were analyzed.

### Protoplast isolation, transfection, and dual-luciferase assay

Protoplast isolation and PEG-Ca^2+^-mediated transfection were adapted from previous study(Yoo et al., 2007). Protoplasts were prepared from leaves of DAI 15 *A. thaliana* plants treated with TRV2-Myc or TRV2-eIF5A, or from *eif5a-ami-es #4* seedlings grown for 5 days after germination and subsequently treated with EtOH or 10 μM Es for an additional 5 days. Tissue was cut into 0.5-1-mm strips, vacuum-infiltrated for 30 min in darkness, and digested for 3 h at room temperature in darkness in enzyme solution containing 20 mM MES-KOH (pH 5.7), 1.5 % (w/v) Cellulase Onozuka R-10 (Yakult Pharmaceutical Industry Co., Ltd., Japan), 0.4 % (w/v) Macerozyme R-10 (Yakult Pharmaceutical Industry Co., Ltd., Japan), 0.4 M mannitol (Sigma-Aldrich, M4125), 20 mM KCl (Sigma-Aldrich, P3911), 10 mM CaCl_2_ (Sigma-Aldrich, C7902), and 0.1 % (w/v) BSA (LPS Solution, BSA100).

The enzyme-protoplast suspension was diluted with an equal volume of W5 solution containing 2 mM MES-KOH (pH 5.7), 154 mM NaCl, 125 mM CaCl_2_, and 5 mM KCl and filtered through a sterile 70-μm nylon Cell Strainer (SPL Life Sciences, 93070). Protoplasts were collected by centrifugation at 100 × g for 2 min, resuspended at 2 × 10^5^ cells mL^-1^ in W5 solution, and rested on ice for 30 min. After removal of the W5 solution, protoplasts were resuspended at 2 × 10^5^ cells mL^-1^ in MMG solution containing 4 mM MES-KOH (pH 5.7), 0.4 M mannitol, and 15 mM MgCl_2_.

For each transfection, 100 μL protoplast suspension containing 2 × 10^4^ protoplasts was mixed with 10 μL plasmid DNA. Each reaction received 5 μg p326-RLUC-(10× codon)-FLUC reporter DNA and 5 μg empty p326-Myc or amiRNA-resistant eIF5A-2-Myc WT, S2A, or S2D plasmid. Protoplasts were transfected with 110 μL PEG-Ca^2+^ solution containing 40 % (w/v) PEG4000 (Fluka, 81240), 0.2 M mannitol, and 100 mM CaCl_2_ for 5 min at room temperature. Transfection was stopped by adding 440 μL W5 solution. Protoplasts were collected at 100 × g for 2 min at room temperature and gently resuspended in 1 mL WI solution containing 4 mM MES-KOH (pH 5.7), 0.5 M mannitol, and 20 mM KCl. Cells were incubated for 16 h at 20-25 °C under a 16-h light/8-h dark photoperiod at approximately 120-150 μmol m^-2^ s^-1^.

Firefly and Renilla luciferase activities were measured sequentially using the Dual-Luciferase Reporter Assay System (Promega, E1910) and a VICTOR5 multilabel plate reader (PerkinElmer) according to the manufacturer’s instructions. Reporter output was calculated as FLUC/RLUC. For *eif5a-ami-es #4* assays, fold change was calculated as (FLUC/RLUC)EtOH/(FLUC/RLUC)ES and normalized to the no-insert control. For VIGS assays, fold change was calculated as (FLUC/RLUC)TRV2-Myc/(FLUC/RLUC)TRV2-eIF5A and normalized to the no-insert control. Each biological replicate consisted of an independently prepared protoplast isolation and transfection experiment.

### Quantitative proteomics following eIF5A depletion

Three independently prepared biological samples of *eif5a-ami-es #4* seedlings were collected for each treatment. Seedlings were grown on MSG medium for 5 d and then transferred to MSG medium containing EtOH or 10 μM Es for an additional 5 d. Total proteins were extracted, quantified by BCA assay(Smith et al., 1985), and digested using a filter-aided sample preparation (FASP) workflow(Wisniewski et al., 2009). Proteins were reduced with 5 mM TCEP for 30 min at 37 °C, alkylated with 50 mM iodoacetamide for 1 h at 25 °C in darkness, washed sequentially with 8 M urea and 50 mM ammonium bicarbonate, and digested with trypsin at an enzyme-to-protein ratio of approximately 1:50 for 18 h at 37 °C. Digestion was stopped with formic acid, and the resulting peptides were desalted using C18 micro spin columns, dried using a SpeedVac, and stored at -20 °C until analysis.

Peptides were analyzed by nano-LC-MS/MS using a UPLC system coupled to a Q Exactive mass spectrometer at eBiogen Inc. Peptides were loaded onto a C18 trapping column (3 μm, 100 Å, 75 μm × 2 cm) and separated on a PepMap RSLC C18 analytical column (2 μm, 100 Å, 75 μm × 50 cm) at a flow rate of 300 nL/min. Mobile phase A consisted of water containing 0.1 % formic acid, and mobile phase B consisted of 80 % acetonitrile containing 0.1 % formic acid. The proportion of mobile phase B was increased from 4 % to 40 % over 120 min, followed by column washing and re-equilibration. MS spectra were acquired over a mass range of 400-2,000 m/z.

Raw data were processed using Proteome Discoverer and searched against the UniProt *A. thaliana* protein database. Acetylation, oxidation, carbamylation, and carbamidomethylation were included in the database search. Protein abundance values were normalized to the total peptide amount, and peptide and protein identifications were filtered at a 1 % false-discovery rate.

### Proteomic data processing and motif analysis

UniProt accessions were mapped to *A. thaliana* Genome Initiative loci using the supplied mapping table, Ensembl Plants BioMart(Bolser et al., 2016; Kinsella et al., 2011), and manual resolution of unmatched isoforms. Normalized abundance values from the three samples per condition were log_2_-transformed. For each protein, log_2_ fold change was calculated as the mean log_2_ abundance in β-estradiol-treated samples minus the mean log_2_ abundance in EtOH-treated samples. Ratio P values exported from Proteome Discoverer were adjusted across proteins by the Benjamini-Hochberg method. Differentially abundant proteins were defined by FDR < 0.01 and |log_2_ fold change| > 1, yielding 60 downregulated and 66 upregulated proteins. Principal-component analysis was performed on centered and scaled log_2_-normalized abundances for 4,607 quantified proteins. For the Figure 5A heatmap, the 60 downregulated proteins were standardized by row to Z scores. Rows were clustered using the default Euclidean distance and complete-linkage method, whereas sample columns were displayed without clustering in the fixed order EtOH1-EtOH3 followed by ES1-ES3. Analyses were performed in R v4.6.1 using tidyverse, ggplot2, ggrepel, pheatmap, ComplexHeatmap(Gu et al., 2016), RColorBrewer, and grid.

TAIR10 peptide sequences (TAIR10_pep_20101214) were assigned based on AGI locus and isoform(Lamesch et al., 2012). Quantified protein groups for which a single sequence could not be resolved, including groups assigned to multiple AGI loci and unmatched isoforms, were excluded. This procedure retained 4,561 sequence-resolved proteins, comprising 60 downregulated, 66 upregulated, and 4,435 background proteins. Sixteen motifs (PPP, PPN, PPD, DPG, PPW, PGE, PPL, PPV, PGD, PPI, WPG, PGA, PPS, PGS, PPE, and PPF) were prespecified based on previously reported eIF5A-sensitive translation contexts(Gutierrez et al., 2013; Schuller et al., 2017) and identified by exact sequence matching. Motif presence was used as the primary endpoint. Overlapping motif-occurrence counts and counts normalized per 100 amino acids were also calculated.

For each motif, a one-sided Fisher’s exact test was used to test enrichment in either the downregulated or upregulated group relative to all other sequence-resolved quantified proteins. P values were adjusted separately across the 16 motifs for each comparison using the Benjamini-Hochberg method. As a length-adjusted sensitivity analysis, binomial logistic regression was performed with motif presence as the response variable and differential-abundance group and protein length as predictors. Odds ratios and 95 % confidence intervals were obtained by exponentiating the corresponding model coefficients. PPW and WPG were omitted from the forest plot because no occurrences were observed, resulting in undefined odds ratios. Analyses were performed using tidyverse, Biostrings, broom, ggplot2, and ggrepel.

### *A. thaliana* eFP Browser analysis

Tissue-expression maps for ATMG00070, AT1G54520, AT3G46660, and AT1G26630 were obtained from the *A. thaliana* eFP Browser(Winter et al., 2007). Images were exported using identical display settings for all four loci.

### Identification of IF5A/eIF5A homologs and conservation analysis

Predicted proteomes were obtained from Ensembl and Ensembl Genomes release 62 as protein FASTA files(Dyer et al., 2025; Yates et al., 2022). *A. thaliana* eIF5A-2 (AT1G26630.1), *E. coli* EF-P (NP_416676.4), and a 132-aa aIF5A sequence from *Thermoproteus tenax Kra 1* (ASM25305v1) were used as competing BLASTP queries against each proteome using BLAST+ with an E-value threshold of 1 × 10^-5^(Camacho et al., 2009). For each target protein, the query hit with the lowest E-value was retained. Candidate family membership was evaluated by RPS-BLAST against the NCBI Conserved Domain Database with an E-value threshold of 1 × 10^-5^(Wang et al., 2023). Eukaryotic eIF5A candidates were required to contain PLN03107 (CDD 215580), whereas archaeal aIF5A candidates were required to contain PRK03999 (CDD 235193). Sequences lacking the expected domain and eukaryotic candidates assigned to the EF-P query were excluded. For loci represented by multiple protein isoforms, one representative sequence was retained by prioritizing the longest sequence, followed by the best CDD and BLAST scores. No additional percent-identity, alignment-coverage, or protein-length threshold was applied. The final curated data set comprised 340 archaeal proteins from 340 species, 657 fungal proteins from 620 species, 587 animal proteins from 484 species, 174 protist proteins from 118 species, and 458 Archaeplastida proteins from 119 species.

Only sequences beginning with methionine were retained for N-terminal residue analysis. For the alignment-free analysis, residues 2-11 were extracted directly from each protein sequence; therefore, the first position in the resulting sequence logo represented the encoded residue immediately following the N-terminal methionine. This analysis was applied to Archaea, Fungi, Protists, Animals, and Archaeplastida. Archaeplastida homologs were additionally aligned using MUSCLE(Edgar, 2004) in MEGA12(Kumar et al., 2024), and the alignment column corresponding to Ser2 of *A. thaliana* eIF5A-2 was identified. Sequence logos were generated using WebLogo 3(Crooks et al., 2004) with identical settings across lineages.

### Taxonomy-based species tree

Species names were mapped to NCBI taxonomy identifiers using ETE3 NCBITaxa(Huerta-Cepas et al., 2016), with manual correction of synonyms where required. The species topology was retrieved using NCBITaxa.get_topology with intermediate nodes disabled and exported in Newick format. NCBI lineage information was used to assign taxonomic ranks. The topology and eIF5A copy-number and clade annotation datasets were visualized in iTOL(Letunic & Bork, 2024). This analysis generated a taxonomy-derived species tree rather than an eIF5A protein phylogeny.

### T-DNA genotyping and molecular validation

Genomic DNA from the DHS and six candidate insertion lines was amplified with gene-specific left- and right-primer pairs and the appropriate T-DNA border primer (LBb or o8409), as listed in Table S1. Homozygous plants were identified by the presence of a border/genic-primer product and the absence of the wild-type left/right-primer product. PCR was performed using PrimeSTAR GXL DNA Polymerase (Takara Bio, R051A). Transcript abundance was measured by qRT-PCR with gene-specific primers and normalized to *PP2AA3*. The DHS lines were retained only for characterization because neither line substantially reduced DHS transcript abundance or hypusinated eIF5A under the tested conditions.

### Structural analysis of eIF5A proteins

Experimentally determined coordinates were obtained from the RCSB Protein Data Bank(Vallat et al., 2026). The crystal structure of *A. thaliana* eIF5A-2 (PDB: 3HKS, chain A)(Teng et al., 2009) and AlphaFold-predicted structures of eIF5A-1 and eIF5A-3(Jumper et al., 2021; Varadi et al., 2024) were analyzed in PyMOL version 2.6.0a0. Non-protein atoms were removed. The eIF5A-1 and eIF5A-3 models were independently aligned to eIF5A-2 by Cα atoms using the PyMOL align command and five outlier-rejection cycles, yielding Cα RMSD values of 1.796 Å over 131 atom pairs and 1.584 Å over 129 atom pairs, respectively. Lys51 was displayed as sticks.

Structures of *Saccharomyces cerevisiae* eIF5A (PDB: 3ER0, chain A) and human eIF5A1 (PDB: 8A0E, chain E)(Wator et al., 2023) were independently aligned to *A. thaliana* eIF5A-2 using the same Cα procedure. The alignments yielded RMSD values of 1.680 Å over 123 Cα atoms and 1.938 Å over 107 Cα atoms, respectively. The terminal NZ atoms of yeast Lys51 and human Lys50 were displaced by 11.45 and 10.65 Å, respectively, from *A. thaliana* Lys51 NZ after superposition.

For comparison with ribosome-bound eIF5A, *A. thaliana* eIF5A-2 was superimposed on hypusinated yeast eIF5A in the 80S complex (PDB: 5GAK)(Schmidt et al., 2016) using all available Cα atoms and the PyMOL super command, yielding a Cα RMSD of 2.102 Å. In the yeast structure, the distance from Hyp51 N1 to P-site tRNA A76 OP1 was 2.7 Å. In the unrefined superposition, *A. thaliana* Lys51 NZ was 0.4 Å from the nearest CCA-end atom, C75 O4’. This sub-angstrom value was treated as steric overlap in an unrefined superposition and not as a physical contact. Structures were rendered with identical orientations, scales, and camera settings within each comparison.

### Statistical analysis

GraphPad Prism v10.5.0 was used for the analyses shown in Figures 1, 3, 4, 5, S1, S5-S7, S9, S12, and S13. Individual data points and definitions of n are provided in the corresponding figure legends. Unless specified otherwise, statistical tests were two-tailed and alpha was 0.05. For primary root length and root hair coverage analyses, outliers were identified and excluded before statistical testing using the ROUT method with Q = 10 %.

EtOH-versus-Es comparisons within genotype in Figures 1C-1D, 3D, 3F, 4B, S1A-S1B, S5E, and S7D were performed using multiple unpaired t tests with Welch correction and Holm-Sidak correction for multiple comparisons. Figure 4D and Figure S7G were analyzed using two-tailed Welch’s t tests. Figures 4G-4H, 5F, and S9 were analyzed by ordinary one-way ANOVA followed by Tukey’s multiple-comparisons test. Figure S6C was analyzed by ordinary one-way ANOVA followed by Dunnett’s multiple-comparisons test against Col-0. Figure S12 was analyzed using multiple unpaired t tests assuming equal SDs with Holm-Sidak correction, and Figure S13 was analyzed using a two-tailed unpaired Student’s t test for each mutant-versus-Col-0 comparison.

Proteomic ratio P values and the 16 motif-enrichment tests were adjusted by the Benjamini-Hochberg method. Motif enrichment was evaluated using one-sided Fisher’s exact tests, and length-adjusted sensitivity analysis used binomial logistic regression as described above. Statistical significance is denoted as ns, P > 0.05; *, P < 0.05; **, P < 0.01; ***, P < 0.001; and ****, P < 0.0001. Exact P values are reported where available.

## Materials availability

All materials generated in this study are available from the corresponding authors upon reasonable request for non-commercial research purposes.

## Funding

This work was supported by the National Research Foundation of Korea (NRF) grants to D.H.L. (RS-2025-25423521) and H.S.L. (RS-2024-00338015), and by the Institute for Basic Science (IBS) grant to H.S.L. and E.Y.K. (IBS-R021-D1-2026-a00). E.Y.K. was additionally supported by the Research Fund for International Scientists (RFIS), National Natural Science Foundation of China (NSFC) (32350610246), the Kunshan Science and Technology Bureau, China (kssc202302072), and the Startup Fund of Duke Kunshan University, China (00AKUG0122).

## Author contributions

**Project Administration**, D.H.L., H.S.P., E.Y.K., and H.S.L.;

**Conceptualization**, D.H.L., C.S.A., H.S.P., E.Y.K., and H.S.L.;

**Investigation**, J.T.Y., D.H.L., S.C., C.S.A., S.M., and Y.L.;

**Supervision**, H.S.P., E.Y.K., and H.S.L.;

**Methodology**, J.T.Y., D.H.L., S.C., and C.S.A.;

**Resources**, D.H.L., C.S.A., H.S.P., E.Y.K., and H.S.L.;

**Formal Analysis**, J.T.Y., D.H.L., and S.C.;

**Validation**, J.T.Y., D.H.L., and S.C.

**Writing-Original Draft**, J.T.Y., D.H.L., and H.S.L.;

**Writing-Review & Editing**, all authors commented on and approved the manuscript;

**Visualization**, J.T.Y., D.H.L., and H.S.L.

## Acknowledgments

We thank all members of the IMGN at Kyung Hee University for valuable discussions and technical support. We are grateful to the plant facility team at IBS for maintaining growth chambers and providing excellent plant care. We also thank the members of CReAtE at Duke Kunshan University for helpful discussions and support throughout this work.

## Declaration of interests

The authors declare no competing financial interests.

---

**Figure S1.**
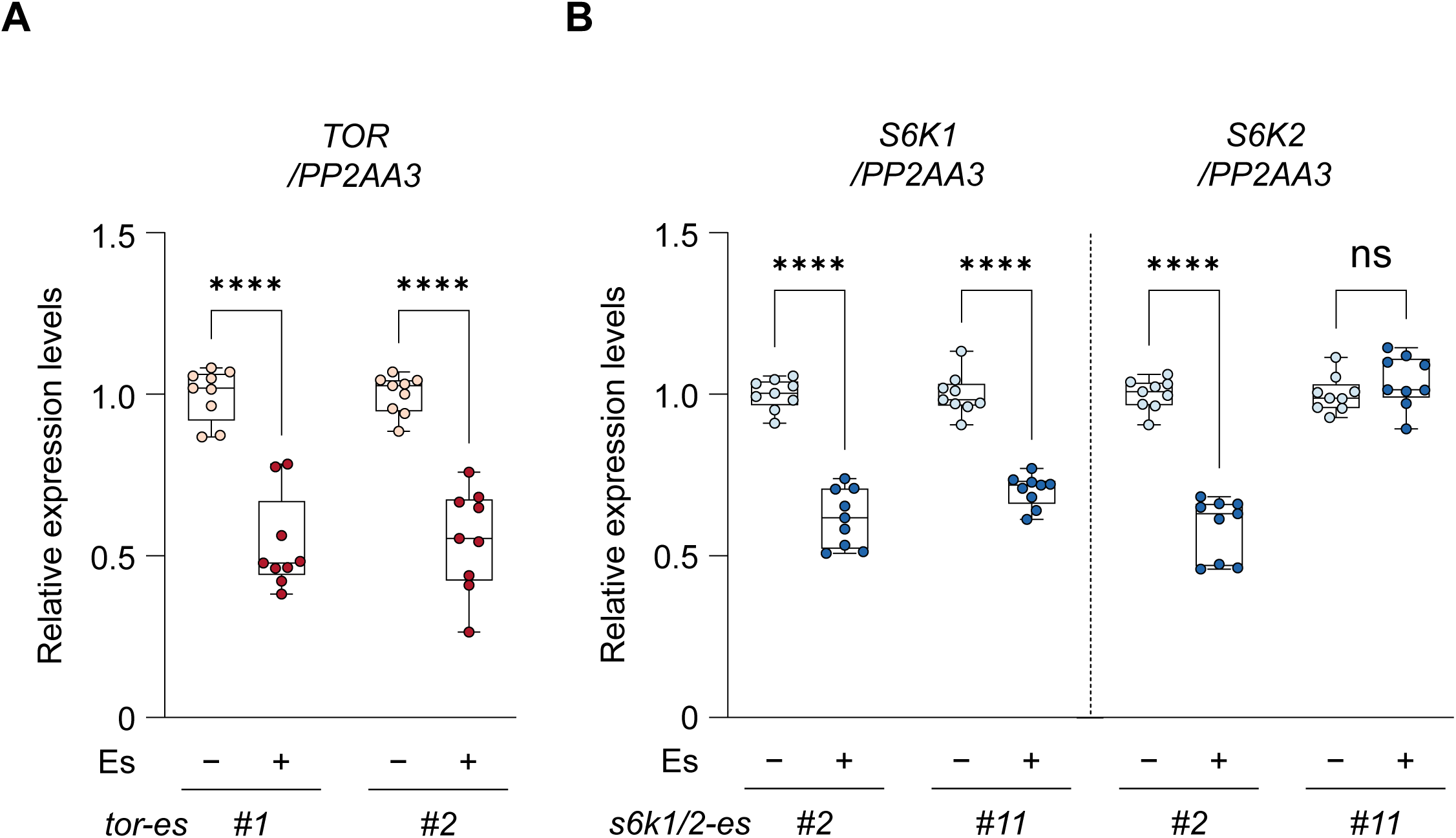
Estradiol treatment reduces TOR and S6K transcript levels in inducible knockdown lines. (A) RT-qPCR analysis of *TOR* transcript levels in *tor-es #1* and *tor-es #2* seedlings treated with EtOH or 10 μM estradiol (Es). Transcript levels were normalized to *PP2AA3*. Box plots show all data points; boxes indicate the first and third quartiles, center lines indicate medians, and whiskers indicate the minimum and maximum values. *n* = 3 biological replicates. Statistical significance was assessed using multiple unpaired t-tests with Welch correction and Holm–Šídák correction for multiple comparisons to compare EtOH and Es within each genotype; ****, P < 0.0001. (B) RT-qPCR analysis of *S6K1* and *S6K2* transcript levels in *s6k1/2-es #2* and *s6k1/2-es #11* seedlings treated with EtOH or 10 μM Es. Transcript levels were normalized to *PP2AA3*. Box plots are shown as in A. *n* = 3 biological replicates. Statistical significance was assessed using multiple unpaired t-tests with Welch correction and Holm–Šídák correction for multiple comparisons to compare EtOH and Es within each genotype; ns, P > 0.05; ****, P < 0.0001.

**Figure S2.**
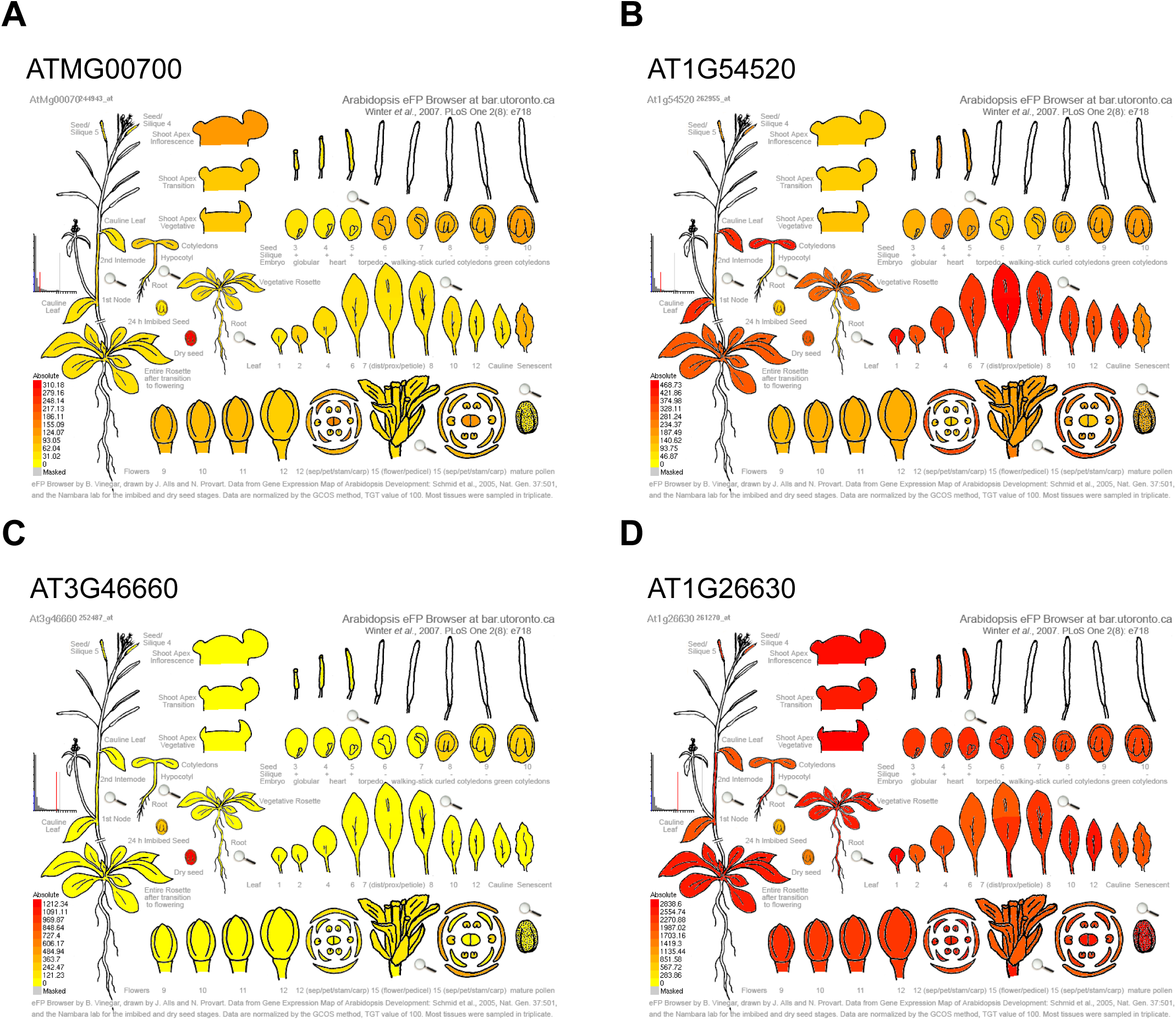
Predicted tissue expression patterns of candidate S6K-associated phosphoprotein genes. (A-D) eFP Browser reports showing the predicted expression patterns of *ATMG00070* (A), *AT1G54520* (B), *AT3G46660* (C), and *AT1G26630* (D) across different *Arabidopsis thaliana* tissues and developmental stages. Expression maps were obtained from the *A. thaliana* eFP Browser. Among these candidates, AT1G26630 shows relatively strong expression in root tissues, supporting its prioritization as a candidate downstream target of S6K relevant to root development.

**Figure S3.**
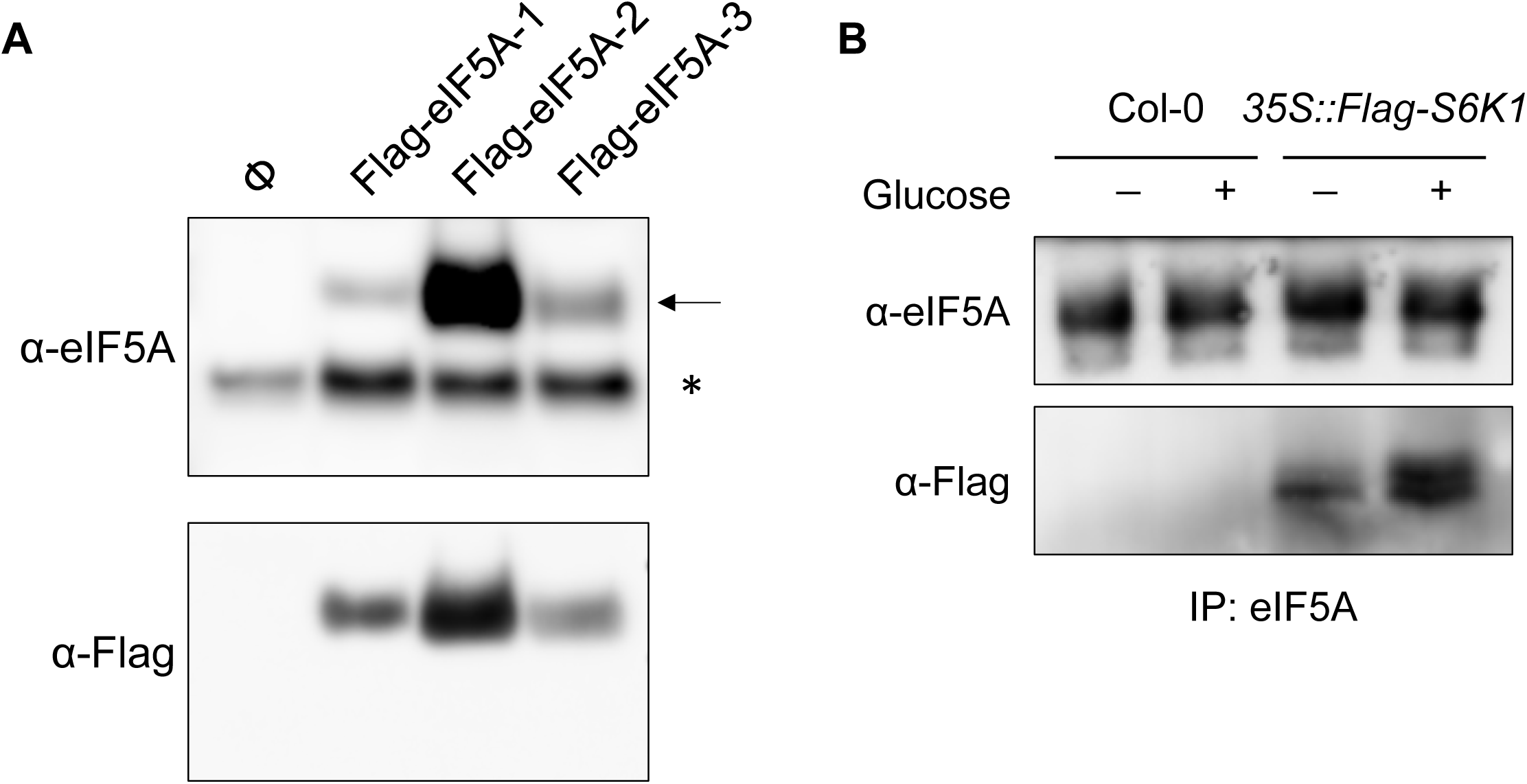
Validation of anti-eIF5A antibody specificity and semi-native S6K1-eIF5A co-immunoprecipitation. (A) Immunoblot analysis showing that the custom anti-eIF5A antibody detects Flag-eIF5A-1, Flag-eIF5A-2, and Flag-eIF5A-3 transiently expressed in *Nicotiana benthamiana* leaves. Anti-Flag immunoblotting confirms expression of the Flag-tagged proteins. The arrow indicates eIF5A, and the asterisk indicates a nonspecific band. (B) Semi-native co-immunoprecipitation of endogenous eIF5A and Flag-S6K1 in *A. thaliana* seedlings. Total proteins from Col-0 or *35S::Flag-S6K1* seedlings treated without or with glucose were immunoprecipitated with anti-eIF5A serum and analyzed by immunoblotting with anti-eIF5A and anti-Flag antibodies. Because input and IP-normalization controls are not shown in this panel, the legend describes detection of co-precipitated Flag-S6K1 without claiming a glucose-dependent increase in interaction strength.

**Figure S4.**
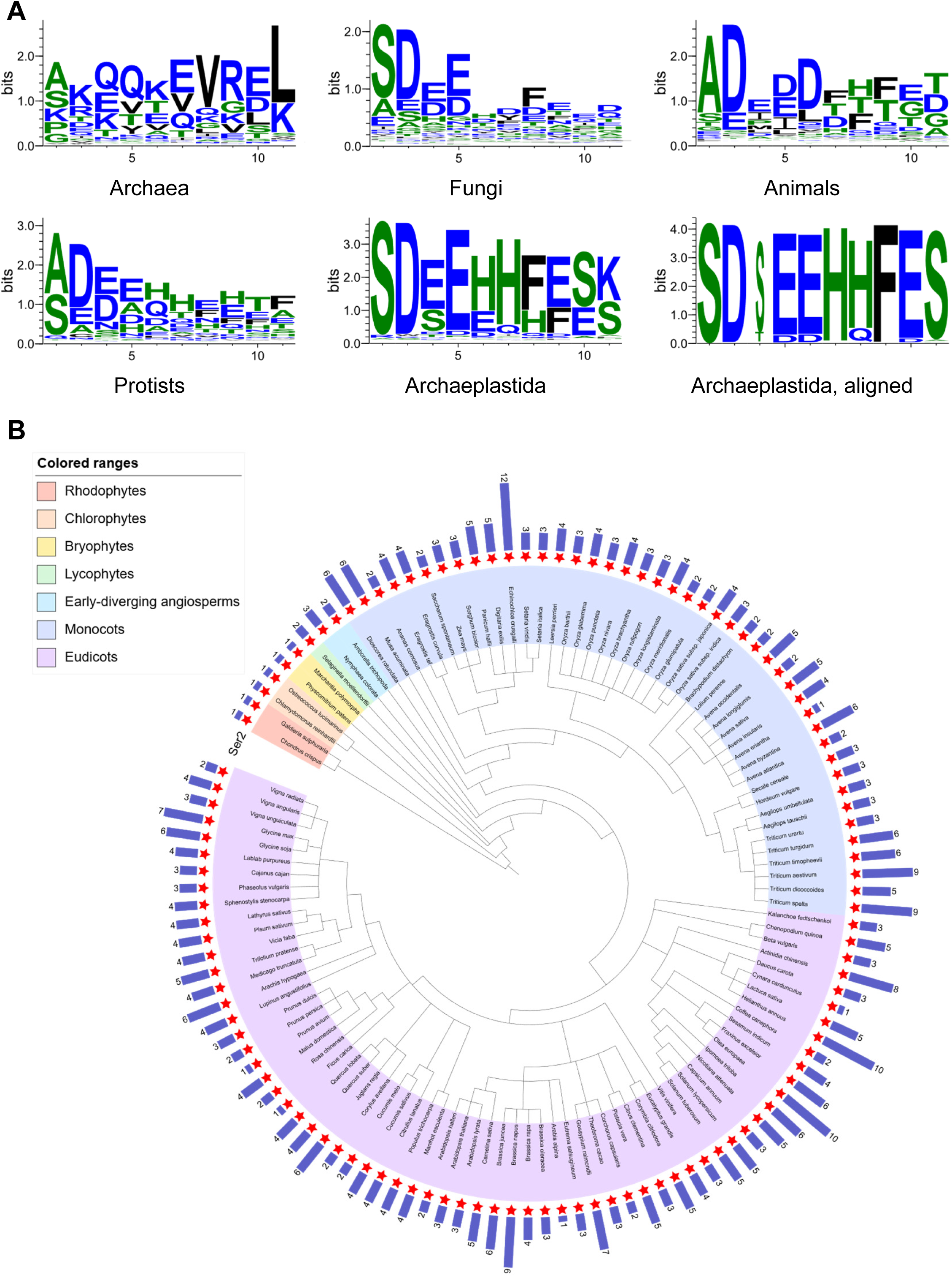
N-terminal residue conservation of IF5A/eIF5A homologs across major lineages. (A) Sequence logos show amino acid frequencies at the N terminus of IF5A/eIF5A homologs from Archaea (340 species, 340 proteins), Fungi (620 species, 657 proteins), Animals (484 species, 587 proteins), Protists (118 species, 174 proteins), and Archaeplastida (119 species, 458 proteins). For Archaea, Fungi, Protists, and Animals, residues were tabulated using an alignment-free approach after removal of the initiator methionine; position 1 in each logo corresponds to the residue immediately following the initiator methionine. For plants, conservation was evaluated both by the same alignment-free approach (“Archaeplastida”) and by Archaeplastida multiple sequence alignment in MEGA12 using MUSCLE to map the residue corresponding to *A. thaliana* eIF5A-2 Ser2 (“Archaeplastida, aligned”). Letter height indicates information content in bits; logos were generated with WebLogo3. (B) Species are arranged according to the taxonomic relationships shown in the innermost circle. The purple-gradient heatmap indicates the number of eIF5A homologs identified in each species. Red stars indicate species in which at least one eIF5A homolog contains serine as the second amino acid residue after initiator methionine removal. Colored ranges indicate major taxonomic clades. This tree represents an NCBI Taxonomy-derived species relationship rather than an eIF5A protein phylogeny.

**Figure S5.**
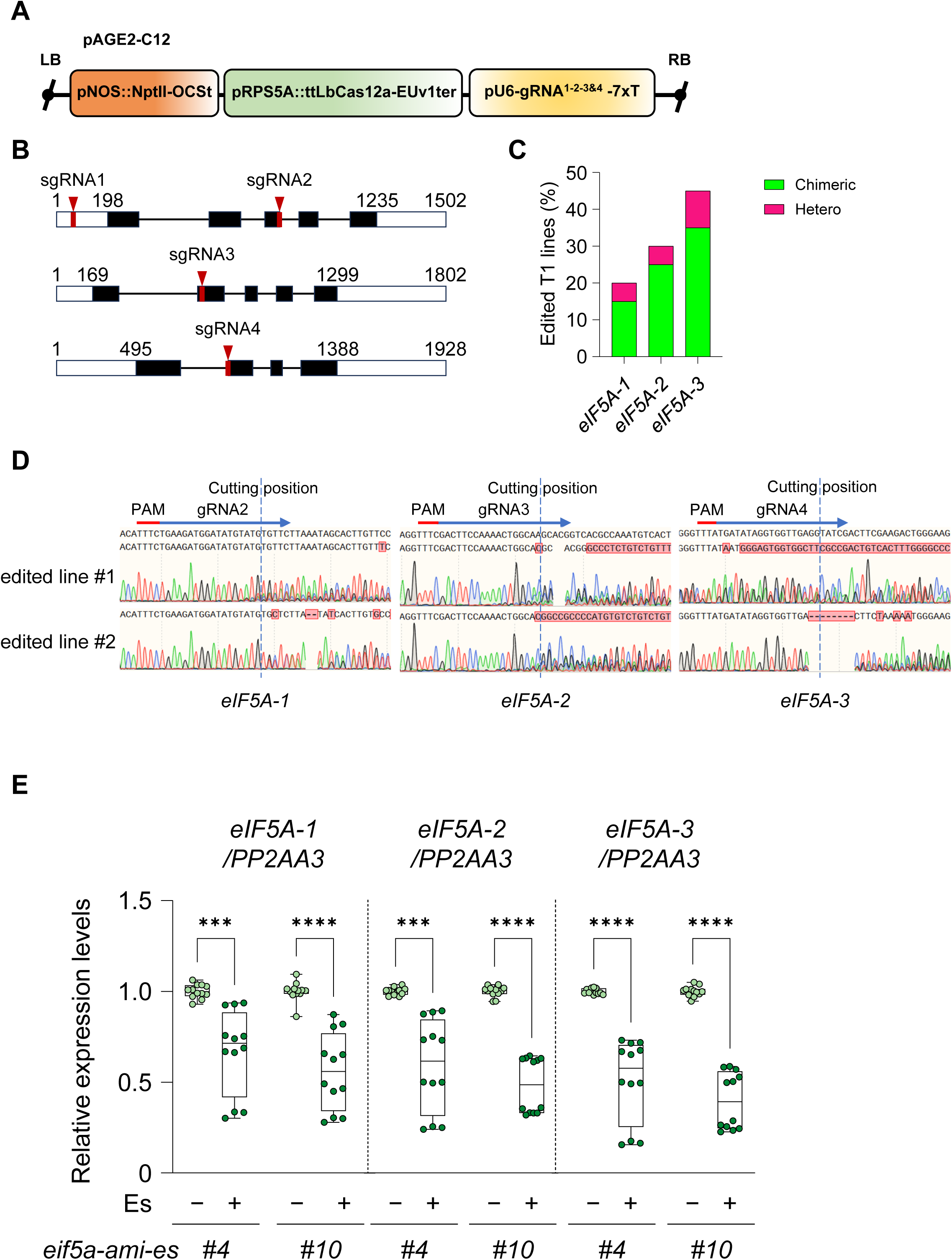
Generation and molecular validation of eIF5A-deficient *A. thaliana* lines. (A) Schematic of the multiplex CRISPR-Cas12a T-DNA construct used to edit the three *A. thaliana eIF5A* genes. The construct contained a pNOS::NPTII-OCSt selection cassette, pRPS5A::ttLbCas12a-EUv1ter, and a U6 promoter-driven array of four CRISPR RNAs (crRNAs). (B) Schematic illustration of the *eIF5A-1*, *eIF5A-2*, and *eIF5A-3* and their corresponding crRNA target sites. White boxes indicate untranslated regions, black boxes indicate coding exons, and connecting lines indicate introns. Red arrowheads indicate the target positions of crRNA1-crRNA4. crRNA1 and crRNA2 target *eIF5A-1*, crRNA3 targets *eIF5A-2*, and crRNA4 targets *eIF5A-3*. Numbers indicate nucleotide positions from the transcription start site. (C) Frequencies of CRISPR-Cas12a editing detected at the three *eIF5A*s among 20 screened T1 plants. Editing outcomes were determined by bulk Sanger sequencing and ICE analysis. Stacked bars indicate the proportions of plants provisionally classified as chimeric/mosaic or putative heterozygous. Editing was detected in 4 of 20 plants for *eIF5A-1* (20 %; three chimeric and one putative heterozygous), 6 of 20 plants for *eIF5A-2* (30 %; five chimeric and one putative heterozygous), and 9 of 20 plants for *eIF5A-3* (45 %; seven chimeric and two putative heterozygous). (D) Representative Sanger sequencing chromatograms of the *eIF5A-1*, *eIF5A-2*, and *eIF5A-3* target regions from WT (Col-0; top) and a representative Cas12a-edited T1 plant carrying edits at all three loci (edited line #1 and #2; bottom). The PAM, corresponding crRNA sequence, and predicted Cas12a cleavage position are indicated. Red boxes, letters, and dashes indicate sequence differences relative to the Col-0 reference. (E) RT-qPCR analysis of *eIF5A-1*, *eIF5A-2*, and *eIF5A-3* transcript levels in *eif5a-ami-es #4* and *eif5a-ami-es #10* seedlings treated with EtOH or 10 μM Es. Transcript levels were normalized to *PP2AA3*. Box plots show all data points; boxes indicate the first and third quartiles, center lines indicate medians, and whiskers indicate the minimum and maximum values. *n* = 4 biological replicates. Statistical significance was assessed using multiple unpaired t-tests with Welch correction and Holm–Šídák correction for multiple comparisons; ***, P < 0.001; ****, P < 0.0001.

**Figure S6.**
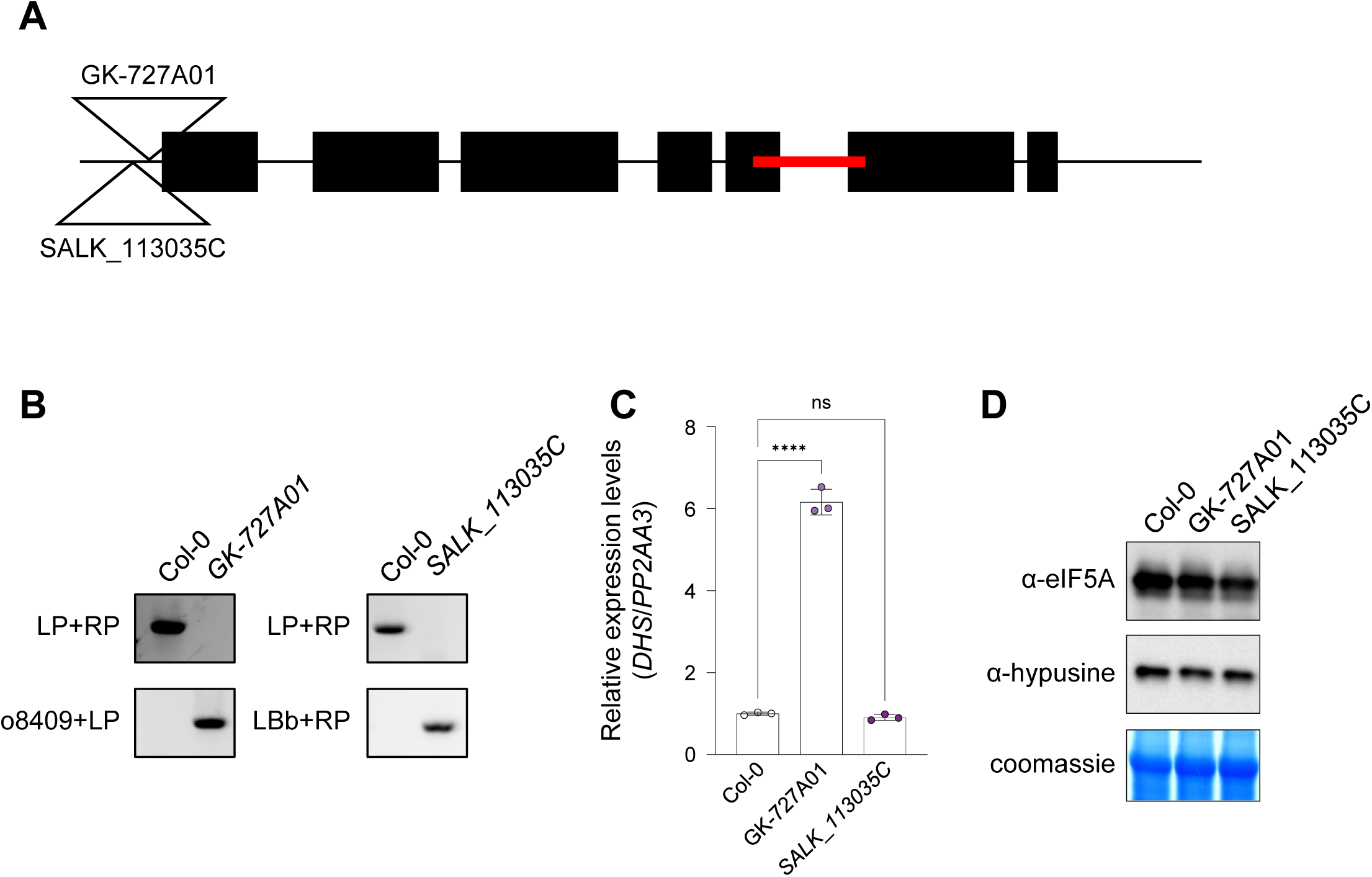
Characterization of DHS T-DNA insertion lines and eIF5A hypusination status. (A) Schematic representation of T-DNA insertion sites in *A. thaliana DHS* mutant lines. Red bars indicate the genomic region amplified for the qRT-PCR assay shown in C. (B) Genotyping PCR analysis confirming homozygous T-DNA insertion lines. (C) RT-qPCR analysis of *DHS* transcript levels in the indicated T-DNA insertion lines. Transcript levels were normalized to *PP2AA3*. Bar graphs show means ± SD. *n* = 3 technical replicates. Statistical significance was assessed using ordinary one-way ANOVA followed by Dunnett’s multiple-comparisons test, comparing each line with the control. ns, P > 0.05; ****, P < 0.0001. (D) Immunoblot analysis of total eIF5A and hypusinated eIF5A levels in the indicated DHS T-DNA insertion lines. CCB staining is shown as a loading control.

**Figure S7.**
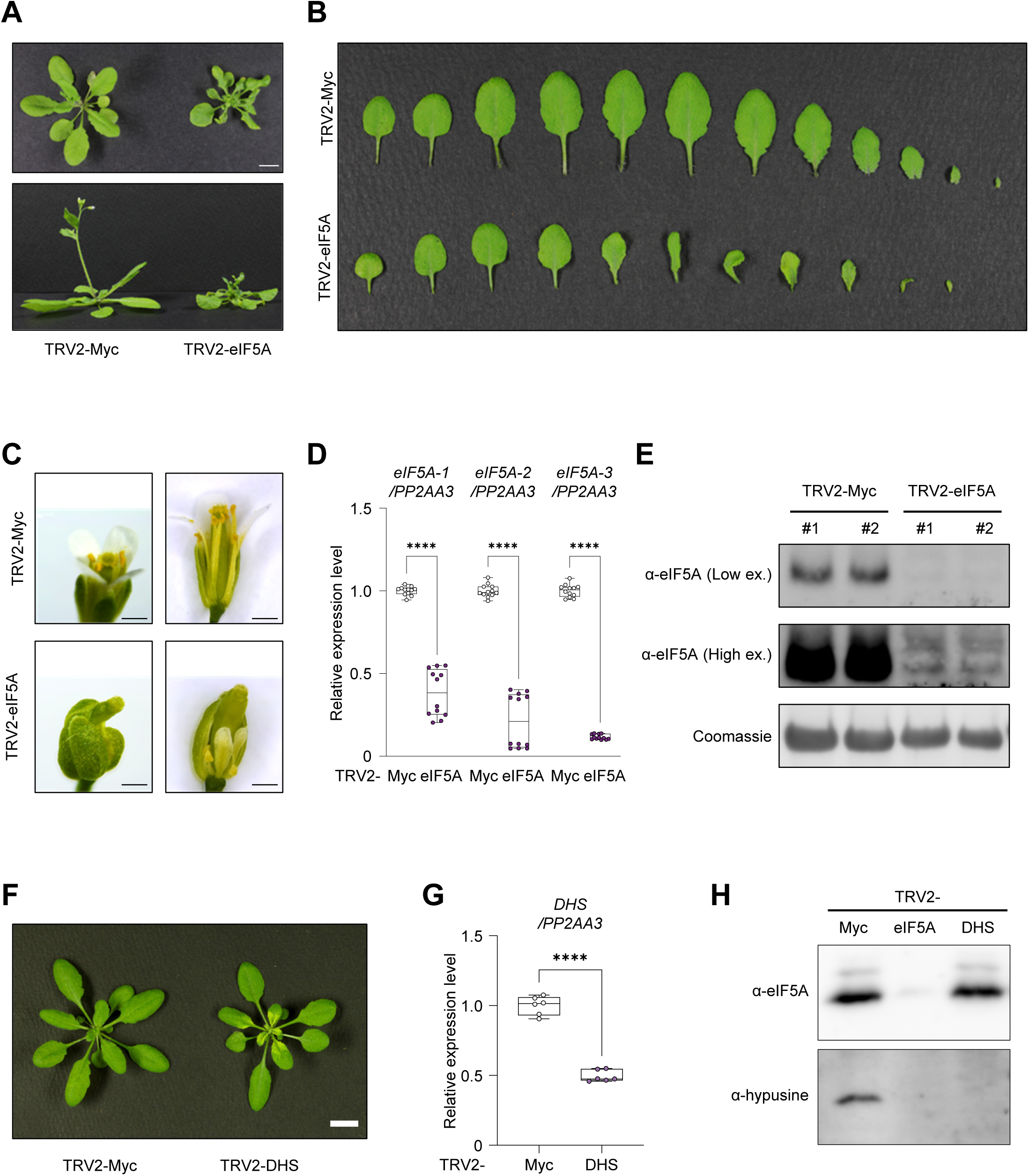
Direct eIF5A silencing causes stronger developmental defects than DHS silencing. (A) Representative vegetative phenotypes of TRV2-myc control and TRV2-eIF5A plants after virus-induced gene silencing (VIGS). (B) Leaf phenotype series of TRV2-myc and TRV2-eIF5A plants. (C) Flower phenotypes of TRV2-myc and TRV2-eIF5A plants. TRV2-eIF5A plants show reproductive abnormalities. (D) RT-qPCR analysis of *eIF5A-1*, *eIF5A-2*, and *eIF5A-3* transcript levels in the ninth leaf at 15 days after infiltration (DAI15) in TRV2-myc and TRV2-eIF5A plants. Transcript levels were normalized to *PP2AA3*. Box plots show all data points; boxes indicate the first and third quartiles, center lines indicate medians, and whiskers indicate the minimum and maximum values. *n* = 4 biological replicates. Statistical significance was assessed using multiple unpaired t-tests with Welch correction and Holm–Šídák correction for multiple comparisons; ****, P < 0.0001. (E) Immunoblot analysis of eIF5A protein abundance in the ninth leaf at DAI15 from TRV2-myc and TRV2-eIF5A plants. Low- and high-exposure anti-eIF5A blots are shown, and Coomassie staining is shown as a loading control. (F) Representative vegetative phenotypes of TRV2-myc and TRV2-DHS plants after VIGS. DHS silencing causes milder visible growth defects than eIF5A silencing under these conditions. (G) RT-qPCR analysis of *DHS* transcript levels in the ninth leaf at DAI15 from TRV2-myc and TRV2-DHS plants. Transcript levels were normalized to *PP2AA3*. Box plots show all data points; boxes indicate the first and third quartiles, center lines indicate medians, and whiskers indicate the minimum and maximum values. *n* = 2 biological replicates. Statistical significance was assessed using a two-tailed Welch’s t-test; ****, P < 0.0001. (H) Immunoblot analysis of total eIF5A and hypusinated eIF5A levels in TRV2-myc, TRV2-eIF5A, and TRV2-DHS plants. *DHS* silencing reduces hypusinated eIF5A without recapitulating the strong developmental defects caused by direct *eIF5A* silencing.

**Figure S8.**
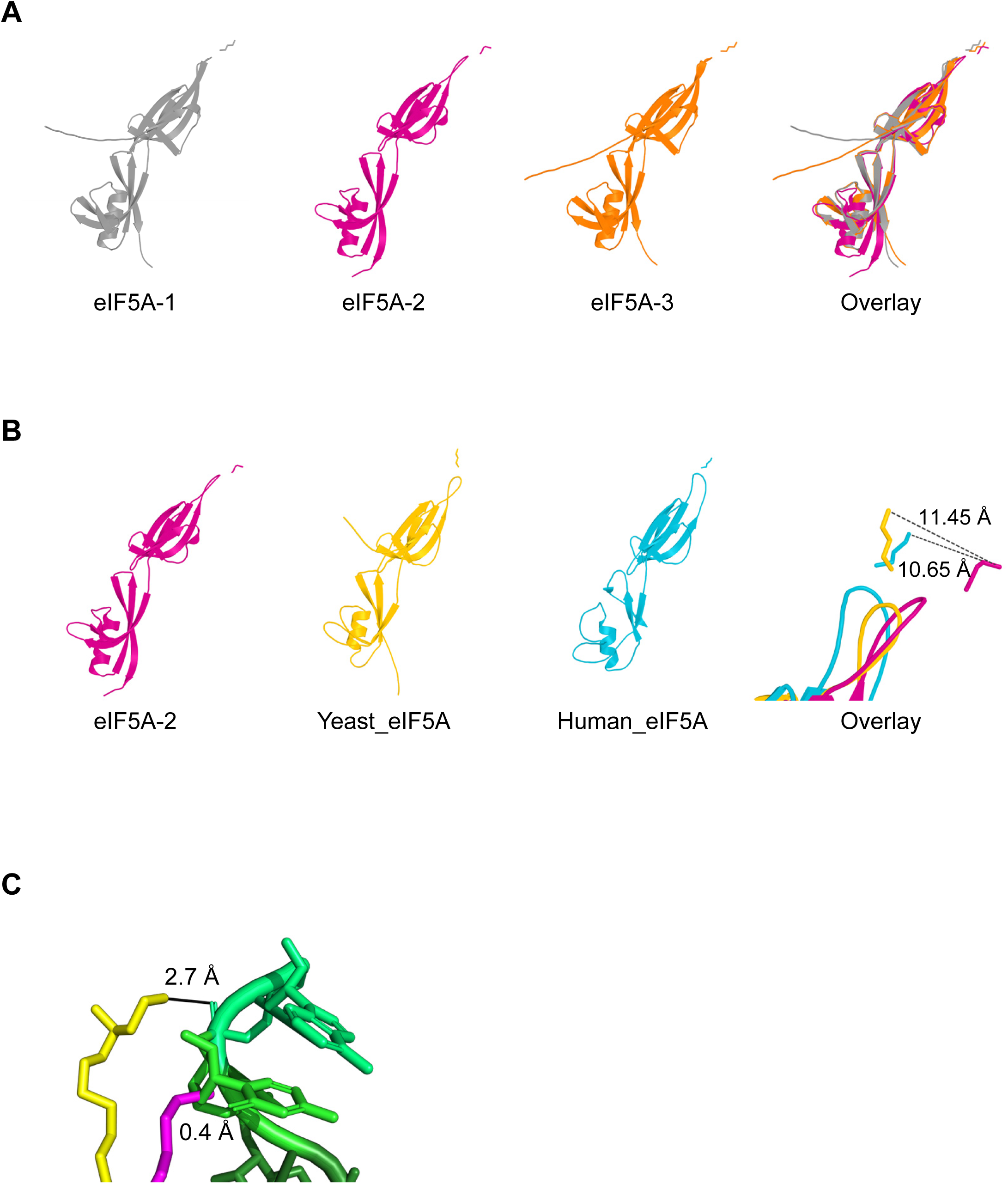
Structural conservation of eIF5A and proximity of the hypusination loop to the ribosomal P-site tRNA. (A) Individual and superimposed structures of *A. thaliana* eIF5A-1 (gray), eIF5A-2 (magenta), and eIF5A-3 (orange) are shown. AlphaFold-predicted structures were used for eIF5A-1 and eIF5A-3 and the crystal structure for eIF5A-2 (PDB: 3HKS, chain A). The hypusination-site residue Lys51 is shown as sticks. Cα-based alignment to eIF5A-2 yielded RMSD values of 1.796 Å for eIF5A-1 and 1.584 Å for eIF5A-3. (B) Structural superposition of *A. thaliana* eIF5A-2 (magenta; PDB: 3HKS, chain A), *Saccharomyces cerevisiae* eIF5A (yellow; PDB: 3ER0, chain A), and human eIF5A1 (cyan; PDB: 8A0E, chain E). Yeast and human eIF5A were independently aligned to *A. thaliana* eIF5A-2, yielding RMSD values of 1.680 Å over 123 Cα atoms and 1.938 Å over 107 Cα atoms, respectively. Lys51 of *A. thaliana* and *S. cerevisiae* eIF5A and Lys50 of human eIF5A1 are shown as sticks. After superposition, the terminal NZ atoms of yeast Lys51 and human Lys50 were displaced by 11.45 and 10.65 Å, respectively, from *A. thaliana* Lys51 NZ. These values describe positional differences among independently determined structures rather than intramolecular distances. (C) Structural superposition of *A. thaliana* eIF5A-2 (magenta; PDB: 3HKS) onto ribosome-bound *S. cerevisiae* eIF5A (yellow; PDB: 5GAK) adjacent to the P-site tRNA CCA end (green). The eIF5A structures were aligned with a core Cα RMSD of 2.102 Å. In the yeast complex, Hyp51 N1 is 2.7 Å from A76 OP1. In the superposed *A. thaliana* structure, unmodified Lys51 NZ is 0.4 Å from the nearest CCA-end atom, C75 O4′. The sub-Å distance reflects steric overlap in the unrefined superposition and should not be interpreted as a physical contact.

**Figure S9.**
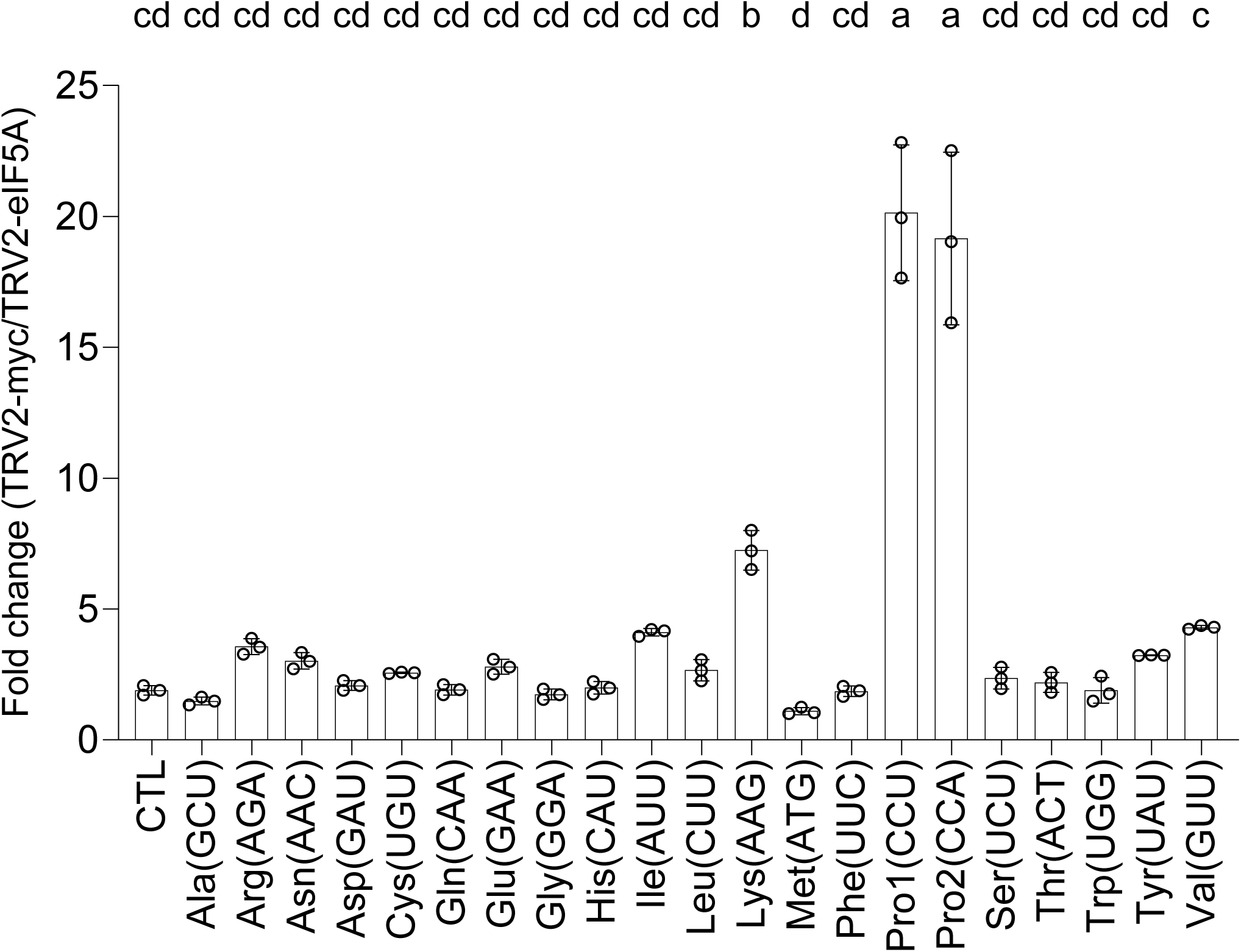
eIF5A VIGS reduces polyproline reporter output. Dual-luciferase reporter assay in protoplasts isolated from TRV2-myc and TRV2-eIF5A plants. Reporters containing 10 repeats of the indicated codon were transfected into protoplasts. Fold change was calculated as (FLUC/RLUC)TRV2-myc/(FLUC/RLUC)TRV2-eIF5A and normalized to the no-insert control (CTL). Bars show means ± SD with individual data points. *n* = 3 technical replicates. Statistical significance was assessed using ordinary one-way ANOVA followed by Tukey’s multiple comparisons test. Different letters indicate significant differences; groups sharing a letter are not significantly different (P < 0.05).

**Figure S10.**
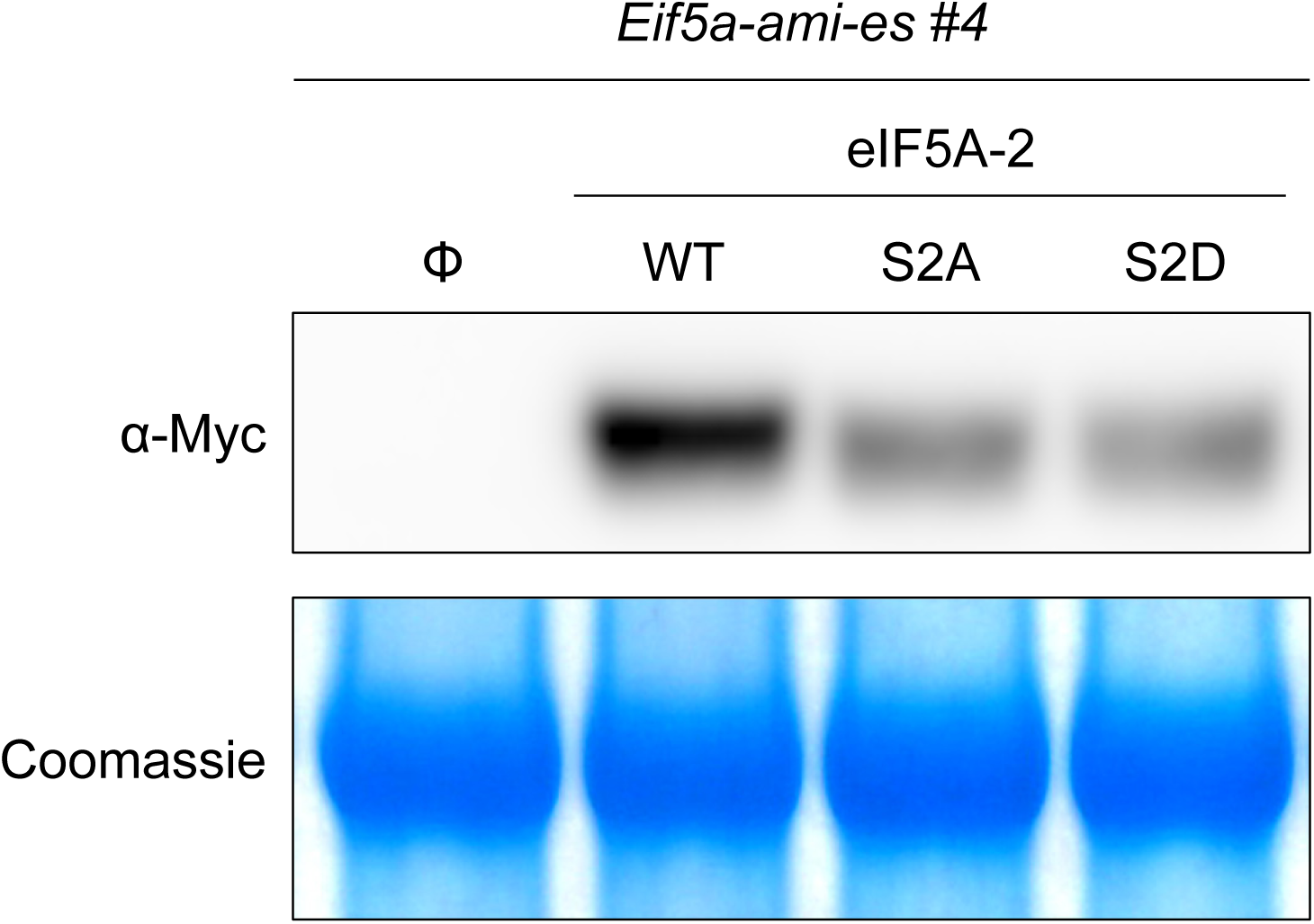
Expression of amiRNA-resistant eIF5A-2 variants in rescue assays. Immunoblot analysis of Myc-tagged amiRNA-resistant eIF5A-2 WT, S2A, and S2D variants expressed in *eif5a-ami-es #4* protoplasts. Φ, empty vector. Proteins were detected using an anti-Myc antibody, and CCB staining is shown as a loading control.

**Figure S11.**
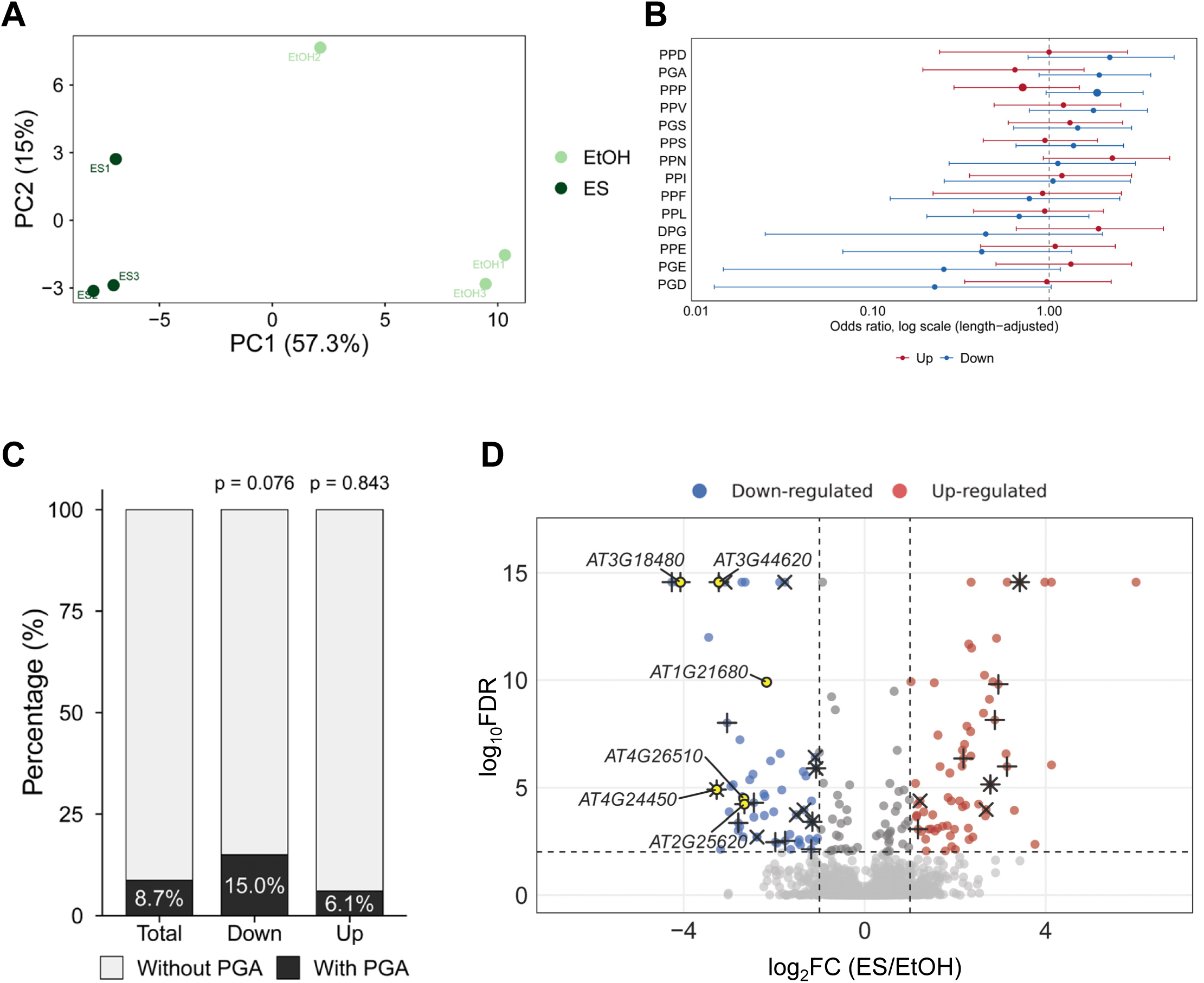
Proteome-wide assessment of proline-rich motifs following eIF5A depletion. (A) Principal component analysis of log_2_-normalized abundances for 4,607 quantified proteins from EtOH- and Es-treated *eif5a-ami-es #4* seedlings (*n* = 3 independently prepared samples per condition). Percentages indicate the variance explained by each component. (B) Length-adjusted odds ratios for motif presence in downregulated (blue) or upregulated (red) proteins. Points show odds ratios from binomial logistic regression adjusted for protein length; horizontal lines show 95 % confidence intervals, and the dashed line indicates an odds ratio of 1. Fourteen motifs with estimable odds ratios are shown. PPW and WPG were excluded because zero observed events yielded undefined odds ratios. The larger PPP points denote the motif tested independently in the reporter assay. No motif reached an FDR of 0.05. (C) Percentages of proteins containing at least one PGA motif among background proteins (labeled Total; *n* = 4,435), downregulated proteins (Down; *n* = 60), and upregulated proteins (Up; *n* = 66). P values were calculated using one-sided Fisher’s exact tests comparing each regulated set with all other sequence-resolved quantified proteins. (D) Volcano plot as in Figure 5C. Black plus and cross symbols indicate significant PPP- and PGA-containing proteins, respectively; yellow points identify the six candidates selected for genetic analysis.

**Figure S12.**
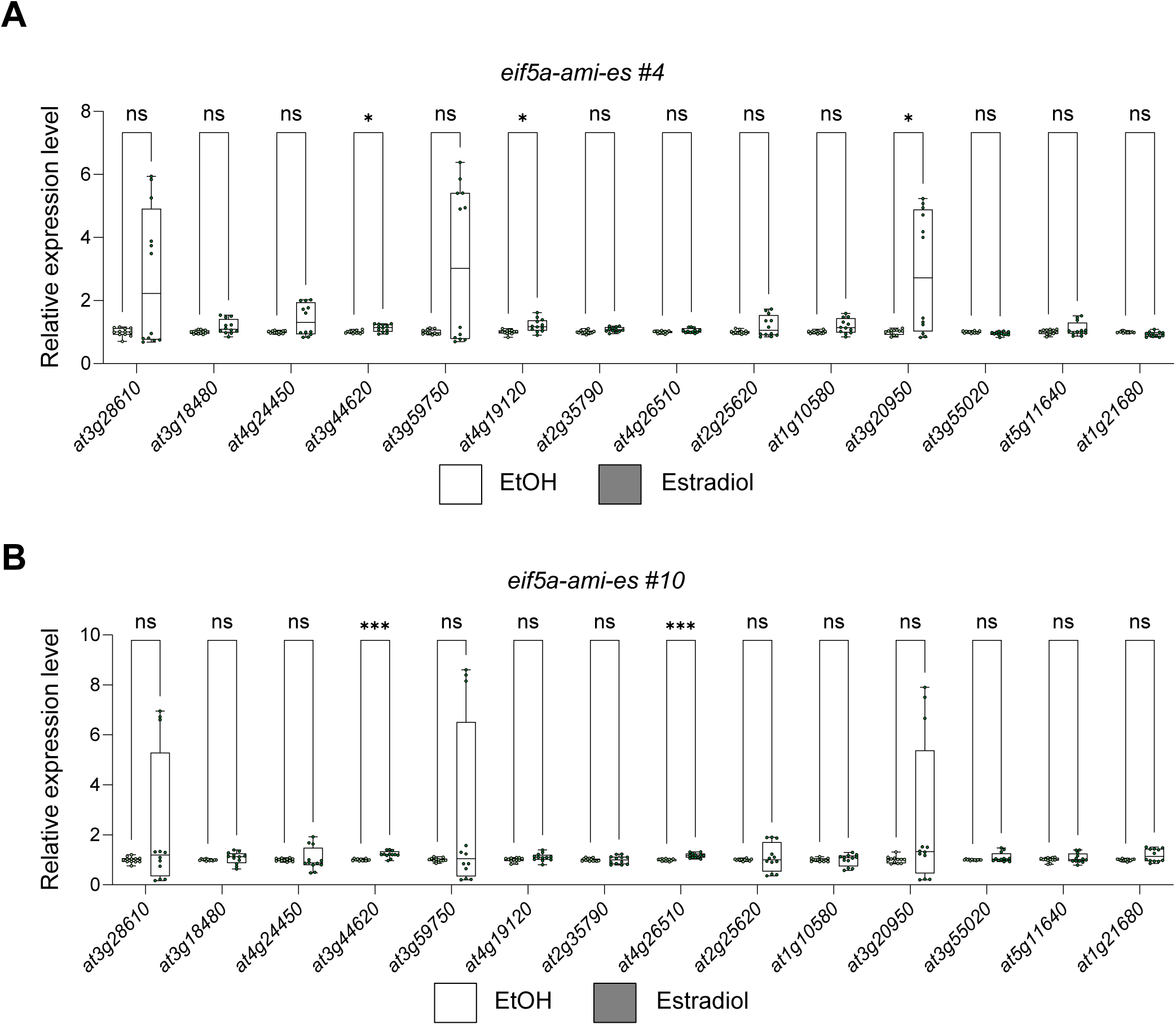
Candidate transcript abundance following inducible eIF5A depletion. (A and B) RT-qPCR analysis of 14 genes encoding proteins reduced at least four-fold in the *eif5a-ami-es #4* (A) and *eif5a-ami-es #10* (B) seedlings treated with EtOH or 10 μM Es. Transcript levels were normalized to *PP2AA3*. Box plots show all data points. *n* = 4 biological replicates. Boxes indicate the first and third quartiles, center lines indicate medians, and whiskers indicate the minimum and maximum values. Statistical significance was assessed using multiple unpaired t tests with the assumption of equal SDs and Holm–Šídák correction for multiple comparisons to compare EtOH and Es within each line; ns, P > 0.05; *, P < 0.05; ***, P < 0.001.

**Figure S13.**
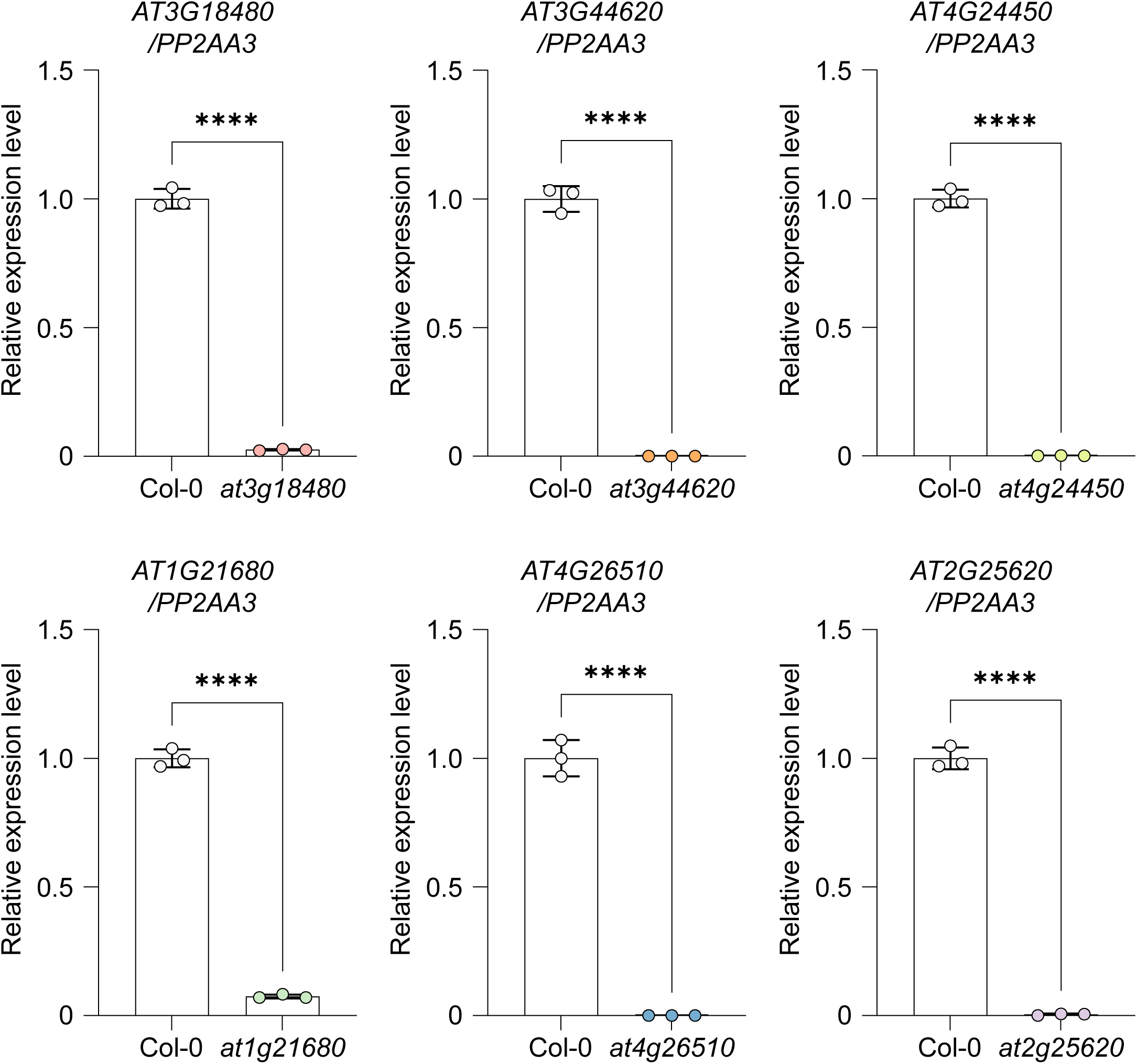
Molecular validation of candidate insertion mutants. RT-qPCR analysis of *AT3G18480*, *AT3G44620*, *AT4G24450*, *AT1G21680*, *AT4G26510*, and *AT2G25620* transcript levels in the corresponding homozygous insertion mutants. Transcript levels were normalized to *PP2AA3* and expressed relative to Col-0. Bar graphs show means ± SD with individual data points. *n* = 3 technical replicates. Statistical significance was assessed using a two-tailed unpaired Student’s t test for each mutant versus Col-0; ****, P < 0.0001.

## Notes

### Competing Interest Statement

The authors have declared no competing interest.

